# Differential active adhesion and a capillary instability drive global eversion and dissemination of bacterial colonies

**DOI:** 10.64898/2026.01.15.699682

**Authors:** Stephan Wimmi, Isabelle Wielert, Kai Zhou, Marc Hennes, Benedikt Sabass, Berenike Maier

## Abstract

Attractive forces between cells determine the shape and sorting behaviour of bacterial colonies. During colony development, chemical gradients form within the colony but it is unclear how they affect cohesion. Here, we discover that such gradients trigger global eversion of proliferating colonies formed by the human pathogen *Neisseria gonorrhoeae*. Like a jet, the inner core flows towards the periphery where it is partially dispersed and partially spreads around the core of the colony. Living dispersed cells reform colonies leading to rapid dissemination. The eversion depends on local oxygen depletion, which reduces the activity of the molecular motors that govern self-attraction: before the eversion the colony consists of a weakly cohesive spherical core surrounded by a strongly cohesive shell. An idealized computational model reveals that in this configuration a non-linear instability occurs, akin to a capillary instability, when the thickness of the strongly interacting shell falls below a critical value. We show that the probability for eversion strongly depends on the motor-generated attractive force between bacteria. Overall, our results indicate that chemical gradients induce spontaneous symmetry breaking in a mechanically active colony, thereby triggering a non-linear shape instability. This instability generates large-scale cellular fluxes that redistribute bacteria toward environments more favourable for their growth.

## Introduction

Biofilms are spatially structured communities formed by microbes. They can protect their inhabitants from external stresses including antibiotic treatment ^1^. Within these biofilms, self-generated chemical gradients cause differentiation into subpopulations with different physiology whose gene expression patterns have been studied for many bacterial species ^2^ ^3^. Since the molecular details differ strongly between species, it is difficult to define unifying principles for biofilm development and dispersal. A promising approach for discovering such unifying principles is to study the physical properties of biofilms ^4^ ^5^ ^6^ ^7^ ^8^ ^9^. The central idea is that, upon coarse-graining, distinct molecular and cellular processes—despite operating far from equilibrium—give rise to convergent mechanical behaviours that ultimately govern biofilm structure and dynamics.

The mechanical properties of bacterial biofilms are governed by steric repulsion, bridging attraction, depletion forces, and osmotic pressure ^7^. The link between adhesin expression and the mechanical properties of bacterial colonies and biofilms has been mainly studied in *Vibrio cholerae* ^10^ ^11^ ^12^, *Pseudomonas aeruginosa* ^13^, and the pathogenic *Neisseriaceae* ^14^ ^15^ ^16^. For rod-like bacteria, such as *V. cholerae* and *P. aeruginosa*, the intrinsic orientational degree of freedom of the elongated cells give rise to nematic order and associated elastic stresses. Thus, growth and ensuing expansion flow couples to orientational ordering in confined environments ^17^, reorientation-driven verticalization enables the transition to three-dimensional architectures ^18^ ^19^, and external flows bias alignment and induce symmetry breaking ^20^. However, a large class of coccoid bacteria, such as *Neisseriaceae*, exhibit far less pronounced shape asymmetry and arguably belong to a different class of active systems.

Here, we will focus on freshly assembled biofilms formed by *Neisseria gonorrhoeae* (gonococcus), henceforth called colonies. The type 4 pilus (T4P) is a ubiquitous adhesin important for cell:cell attraction in various bacterial species ^21^. The fact that the T4P is a molecular motor that continuously consumes energy to elongate and retract ^22^, adds an interesting level of complexity to the system: each bacterium in the colony undergoes a continuous tug-of-war within colonies. The resulting motility correlates with T4P density, T4P motor activity, and T4P:T4P binding rate ^23^ ^24^ ^15^ ^25^. The attractive forces between the cells at the microscopic level govern the surface tension and viscosity of the colony and give rise to a spherical colony shape ^15^ ^14^.

The differential strength of adhesion hypothesis predicts that cells exerting differential attractive forces on each other will segregate ^26^. Extracellular molecular motors used for cellular adhesion result in a tug-of-war between neighbouring cells, whereby the cell bodies move in the direction of the strongest, least breakable bonds. Although the differential strength of adhesion hypothesis was originally developed for understanding cell sorting during embryonic development ^26^, it applies to gonococci that use their T4P for actively pulling on neighbouring cells ^27^ ^28^. When gonococci with different T4P activities are mixed, the cells with lowest attractive force sort to the periphery of the colony ^27^ ^29^ ^30^ ^28^ ^16^ ^31^ ^23^. While similar sorting principles were found also for other bacterial species ^32^ ^33^, it is unclear how the differential strength of adhesion principle affects the structure and dynamics of developing colonies where attractive forces may change with time due to local cellular differentiation. Here, we discover a previously unrecognised emergent phenomenon. Namely, a global eversion of bacterial colonies, in which cells from the colony centre are transported toward the exterior. By combining single-cell tracking in three-dimensional colonies with simulations, we show that this eversion is driven by a reduction in attractive pili-mediated interactions at the colony centre, arising as a consequence of oxygen depletion. This eversion process drives removal of nonviable cells from the colony core and promotes cell dissemination.

## Results

### Global colony eversion drives live and dead cells to the periphery

To find out how the physical properties of a bacterial aggregate develop as the newly formed aggregate grows into a mature biofilm, we cultivated gonococcal colonies of *N. gonorrhoeae* under constant nutrient supply in a microscopic flow chamber. We followed their maturation over time by simultaneously examining the morphology of the colonies in brightfield and staining extracellular DNA (eDNA) and dead cells with SytoX (Fig. 1). The colonies were spherical in shape and their radii increased over time due to cell proliferation and colony fusion in the first 1 - 5 h (Fig. 1A, Extended Data Figures 1-2, Movie S1-3). During the process, eDNA and dead cells accumulated at the centre of the colonies.

**Fig. 1.**
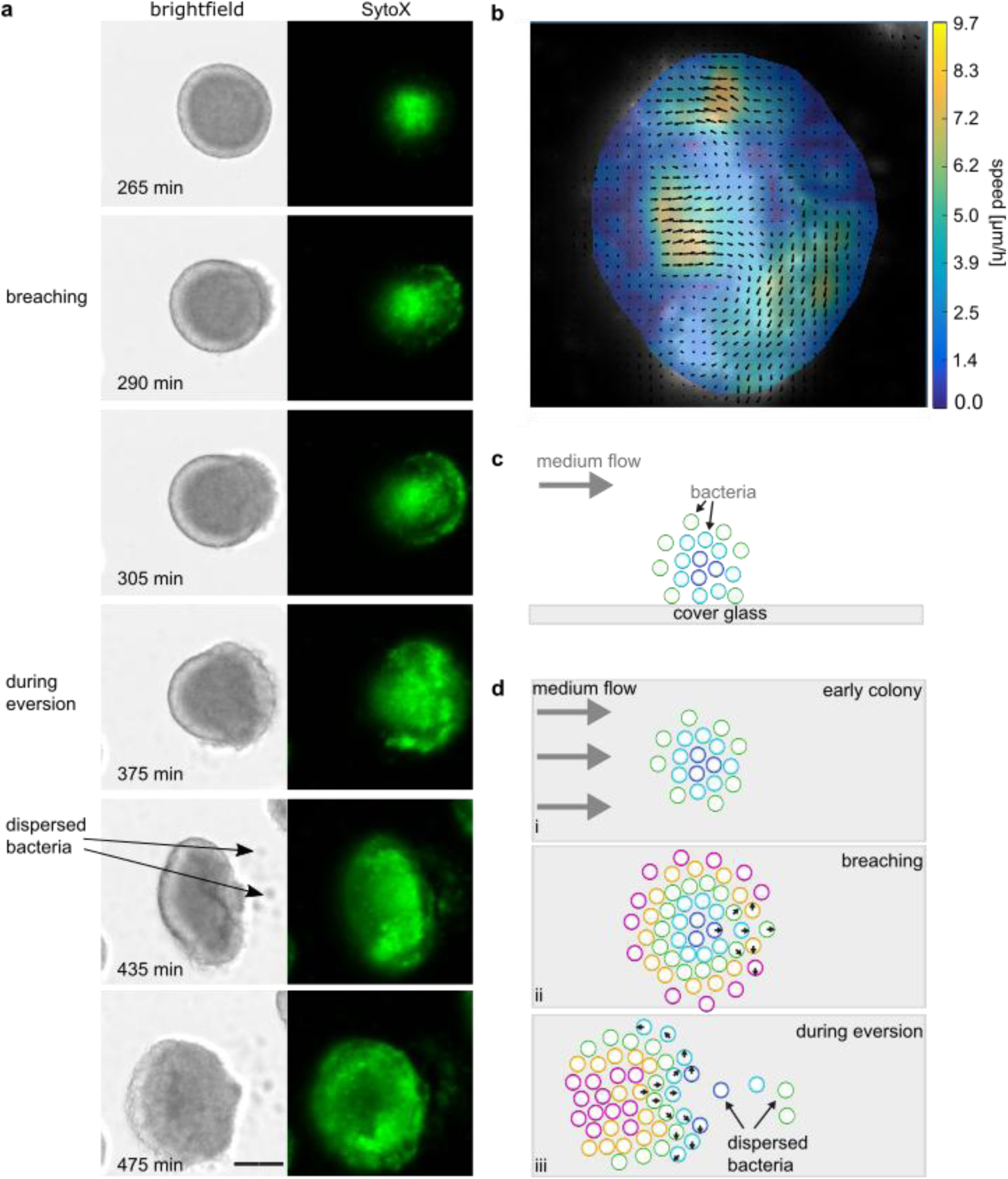
Colony eversion generates ballistic flow from colony centre to periphery. **a)** Time lapse of *N. gonorrhoeae* (NG150) colony incubated in flow chamber in brightfield (left) and SytoX fluorescence mode (right). In each image, the SytoX signal was normalized to 0.2% pixel saturation. Scale bar: 40 µm. Comparison to equalized micrographs is shown in Extended Data Figure 1. This behaviour was observed in 7 biological replicates with N = 30 colonies. **b)** Particle image velocimetry (PIV) analysis of SytoX signal corresponding to time lapse shown in a) over 100 min. **c)** Schematic side view of a colony growing on a surface. **d)** Schematic time lapse of growing colony (top view). i) Early colonies have a spherical shape. ii) At the start of the eversion, cells start moving ballistically and breach the surface. iii) During eversion, formerly central cells (cold colours) spread around the former edge. A subpopulation of cells is dispersed. Colours indicate the original position of cells within the colony. Arrows indicate the net direction of movement during colony eversion.

This phenotype changed strikingly at ∼ 5 h of incubation and at an average radius of 23 µm (Fig. 1, Extended Data Figures 1-3, Movie S1-3). We observed a directional flow of cells from the centre of the colony to its edge and a breach in the surface of the spherical colony. Most cells that have reached the edge spread around the original surface. At this point, the entire microcolony turned inside out, performing a complete eversion (Fig. 1). Henceforth, we will refer to this phenomenon as eversion. Using particle image velocimetry (PIV), we determined the spatiotemporal dynamics of dead cells and extracellular DNA stained by SytoX (Fig. 1b, Extended Data Figure 2). Dead cells and eDNA accumulated at the centre as the colonies grew (Fig. 1a, Extended Data Figures 1-2, Movie S4). During the eversion process, the dead cells flowed towards the periphery in a directed stream and breached the outer shell of the colony. A part of the outwards flowing biomass was redirected over the surface of the colony and stayed attached to the new periphery of the inverted colonies (Fig. 1d). The entire process took several hours and the direction of the eversion was random relative to the direction of medium flow (Extended Data Figure 3c). The fraction of colonies starting eversion increased as a function of colony radius (Extended Data Figure 3d), indicating that colony size affects eversion probability.

Overall, we discovered that gonococcal colonies evaginate during maturation, everting the existing colony inside out.

### Local oxygen depletion causes a reduction of attractive forces at the colony centre

To explain the massive re-organisation of the colony, we investigated the spatio-temporal dynamics of attractive forces between neighbouring bacteria. We expected that the cohesive forces are homogeneous throughout newly assembled colonies. As the gonococcal colony matures, radial oxygen gradients form within the colonies, correlating with reduced membrane potential ^5^. Since the depletion of proton motive force reduces pilus density ^34^, we hypothesise that reduced oxygen concentration at the centre of the colony might affect the T4P adhesin density (Fig. 2a). If T4P density was reduced, cells with weaker attractive force would reside in the centre.

**Fig 2.**
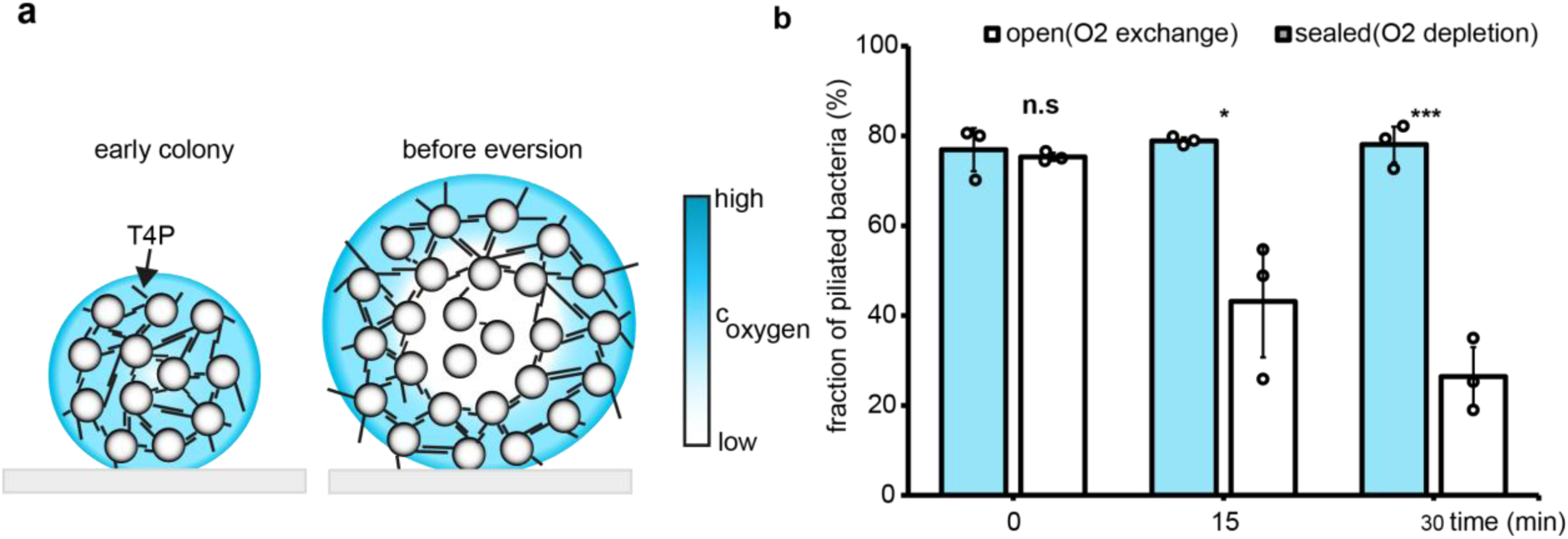
Oxygen depletion reduces piliation. **a)** Sketch of suggested model for gradient formation in gonococcal colonies. T4P-mediated cohesion causes the formation of nearly spherical colonies. Due to respiration, bacteria generate an oxygen gradient within the colony. As a consequence, the number of T4P per cell is reduced at the centre of the colony. **b)** Quantification of single cells displaying T4P over time with air-exchange (white) in an air-tight chamber (gray). *N. gonorrhoeae* expressing a cysteine mutant of the major pilin subunit (*pilE_cys_,* NG226) covalently labeled with Alexa Fluor 488 C5 maleimide was imaged. N = 82-112 cells per condition, and from 3 independent replicates have been analysed. Small circles indicate individual biological replicates, and whiskers denote standard deviation. n.s.: 0.6666; *: p<0.1; ***: p<0.001.

To test this hypothesis, we first verified that the eversion process is dependent on T4P motor dynamics. The eversion experiment was performed with a *pilT* deletion strain which lacks the T4P retraction motor PilT but still forms colonies ^35^. In the flow chamber, the colonies grew, but over a time period of ∼20 h, no eversion, major re-organisation, or outward flow of cells could be observed (Extended Data Figure 4, Movie S5). This result indicates that T4P retraction activity is essential for colony eversion.

Next, we investigated the effect of oxygen depletion on T4P density, by directly visualizing single T4P during incubation in oxygen-permeable and oxygen-impermeable chambers. We found that bacteria rapidly lost piliation and motility in oxygen-impermeable chambers (Fig. 2b, Extended Data Figure 5, Movie S6) consistent with reduced attractive force ^36^.

If oxygen depletion triggers colony eversion, then reduced oxygen levels in the flow chamber should result in earlier eversion events. To create such conditions, we reduced the flow speed during the eversion experiment (from 2.5 rpm to 0.5 rpm). As expected, the changed conditions resulted in earlier eversions (Extended Data Figure 6, Movie S7, 8). Even though the duration of the eversion process was shorter and the morphological changes were less pronounced, we observed directed flow of eDNA from the centre to the periphery as before, further supporting the conclusion that depletion of oxygen triggers eversion.

Finally, we addressed the question whether denitrification prevents colony eversion. To thrive within the human body and under oxygen-limiting conditions, *N. gonorrhoeae* is capable of switching from aerobic respiration to denitrification. We have shown previously, that denitrification enables gonococci to maintain electrical polarization ^5^ ^37^ and expect that oxygen is not required for maintaining piliation. To assess this point, we added 5 mM sodium nitrite (NaNO_2_) to the medium of the eversion experiment. Strikingly, the presence of NaNO_2_ prevented the eversion completely (Extended Data Figure 7). Although small colonies displayed growth arrest after 3-4 h incubation within NaNO_2_ - supplemented media, a subset of larger colonies (∼ 12 %) grew three times bigger than under standard conditions, up to the diameter of ∼150 µm, over a maximal duration of 17 h without displaying any signs of eversion. Taken together, our data strongly point to a scenario, in which oxygen consumption and colony growth form an outside-in oxygen gradient that results in a reduced piliation of cells at the colony centre triggering eversion.

Next, we intended to visualize the spatio-temporal dynamics of attractive interactions between neighbouring cells in developing colonies. We have shown previously that the motility of single cells decreases as a function of increased piliation and, hence, attractive force ^24^ ^38^ ^34^. Thus, we monitored single cell motility with spatial resolution in maturing colonies by acquiring the trajectories of single cells expressing cytosolic *gfp*. Within newly assembled colonies bacteria are fairly immobile and only cells residing within a few micrometres from the edge of the colony show strong motility (Fig. 3a)^24^ ^38^ ^25^. We attribute the strong surface motility to reduced cell:cell interactions and will ignore this highly motile surface layer in the following. In the centre of young colonies (2-3 h before eversion), analysis of the mean-squared displacement ⟨Δx^2^⟩ ∝ t^α^ revealed a sub-diffusive motion with α= 0.3 (Extended Data Figure 8) consistent with caged dynamics ^24^. One hour prior to eversion, the scaling exponent increases to α= 0.6, and during eversion, the motility is characterized by α= 1.4.

**Fig. 3.**
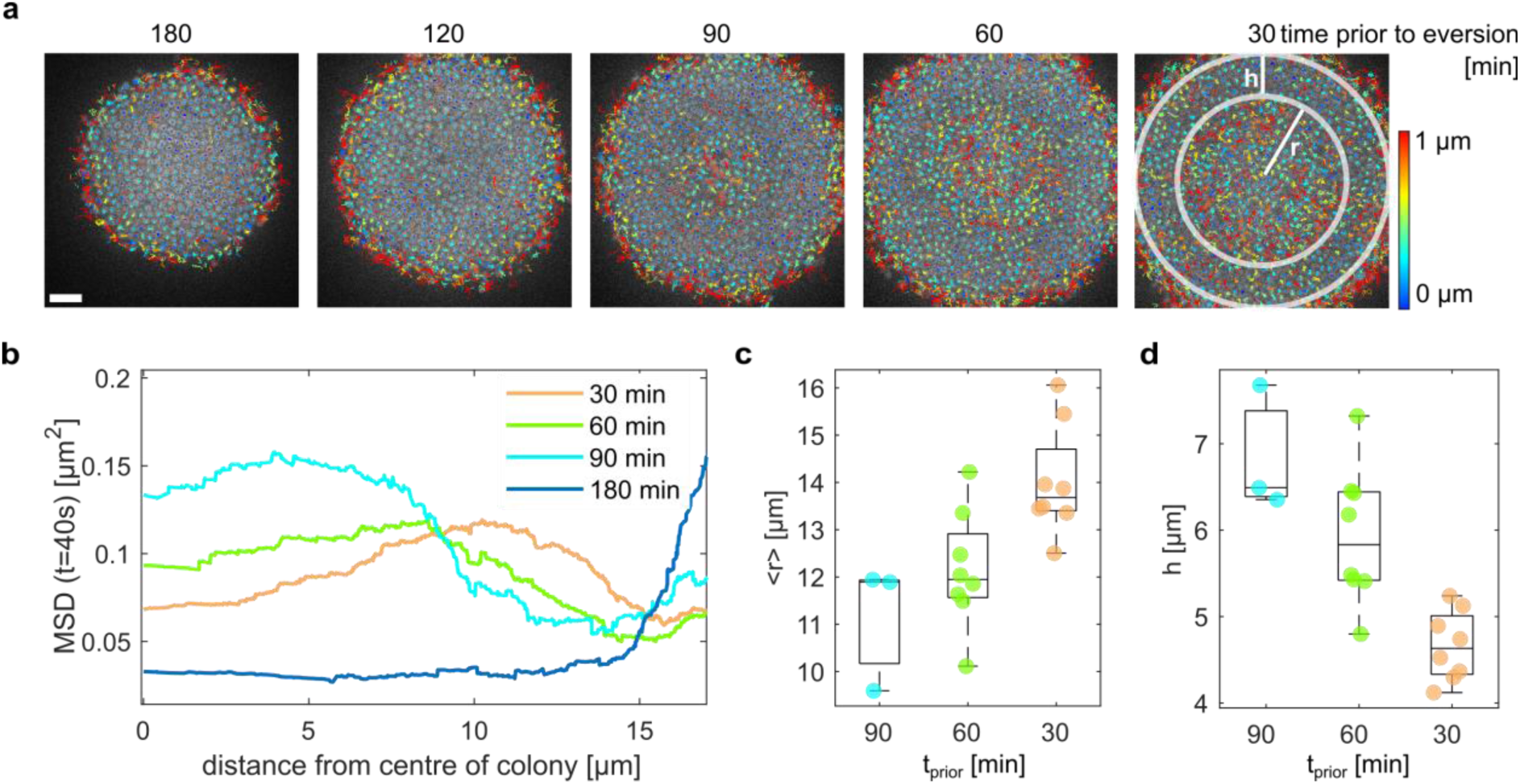
Single cell motility indicates reduction of attractive force at the colony centre. **a)** Overlay between GFP fluorescence and single cell trajectories over 4 min at different times prior to colony eversion. Scale bar: 5 µm. Colour legend: track displacement. Strain: NG151. **b)** Example of mean-squared displacement of single cells after 40 s versus distance from the centre of the colony. The hypermotile surface is removed for clarity. Additional examples, including the surface, are shown in Extended Data Figure 9. **c)** Radius r of hypermotile layer as a function of time prior to colony eversion. blue: 120 min, cyan: 90 min, green: 60 min, orange: 30 min prior to eversion. **d**) Thickness h of the layer of strongly interacting bacteria. Box plots show the median (central mark), bottom and top edges of the box present 25^th^ and 75^th^ percentiles of the data, respectively. N = 8 colonies from 5 biologically independent experiments.

We next characterized single cell motility throughout the entire colony. The magnitude of movement was quantified by calculating the mean-squared displacement of single cells within a time period of forty seconds. Between two and one hours before the start of colony eversion, single cell motility increased at the centre of the colony, indicating that T4P-mediated attractive forces were reduced as a consequence of oxygen limitation (Fig. 3a, b). The hypermotility at the centre transformed into a shell of hypermotile cells that travelled radially through the colony (Fig. 3b, c, Extended Data Figure 9, Movie S9). As a consequence, a shell of cells with low mobility, i.e. strong attraction, surrounded a core of hypermotile cells. The thickness of this strongly attractive shell, h, decreased as a function of time (Fig. 3d, Extended Data Figure 9).

In summary, oxygen depletion reduces T4P density and thus retractile activity. In agreement with this finding, a shell of hypermotile cells travels radially through the colony before the eversion. Since increased motility indicates decreased cellular attractions, the colony comprises a core of weakly attractive cells surrounded by a shell of strongly attractive cells. By contrast, the differential adhesion hypothesis would suggest that the system assumes a minimum-energy state if the strongly attractive cells segregate to the centre of the colony. In the next section, we will assess whether this prediction from equilibrium physics is also found to be true for active bacterial colonies.

### A non-linear shape instability triggers remodelling of the colony

Gonococcal colonies are nearly spherical, and colony eversion represents a spontaneous, violent symmetry breaking. To understand the physics of colony eversion, we next investigated the colony dynamics in an in-silico model, simulating the stochastic dynamics of bacteria and T4P interactions. Experimentally, we have shown that after 4 h of colony growth, the growth rate of bacteria at the centre of the colony is very low ^39^. In simulations, we examined how various initial configurations can drive colony eversion. Starting with a configuration consisting of a shell of strongly attractive cells with active T4P surrounding a core of bacteria with reduced T4P activity (Fig. 4a), we observed two qualitatively different types of behaviours. Firstly, we find biased diffusive motion of less active cells towards the periphery. This diffusive mixing eventually leads to the reverse configuration with the active, contractile bacteria at the core. However, secondly, we also observed a mechanical instability that appeared as a large-scale inside-out eversion, pushing the core in a jet-like fashion to the periphery, (Fig. 4a, b, and Movie S12). This eversion required active retraction of T4P in simulations. A proportional reduction of T4P activity in the outer and inner part of the colony reduced the probability for eversion, see Extended Data Figure 12. Thus, the non-equilibrium conditions set by bacterial activity are required for observing the instability in a parameter range appropriate for modelling bacterial colonies.

**Fig. 4.**
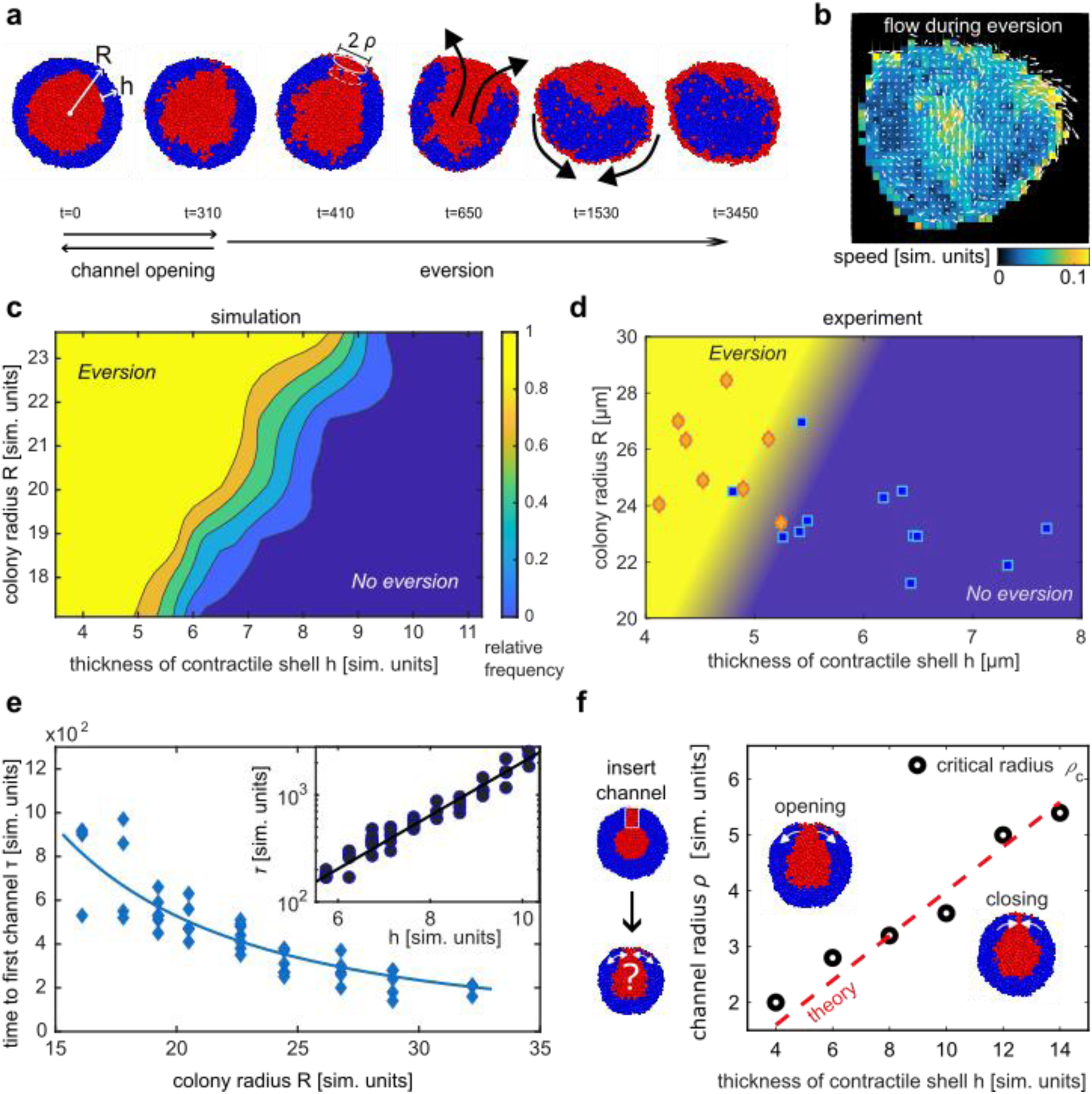
Simulations reveal a non-linear capillary instability. **a)** Simulation snapshots of sections through a colony consisting of two types of cells, undergoing eversion. Initially, bacteria with T4P form a contractile shell (blue) around a core of bacteria with reduced T4P activity (red). The process consists of an initial reversible phase, during which passive bacteria form a channel in the contractile shell, followed by a large-scale deformation of the colony. **b)** Averaged cell motion during the eversion. **c)** Simulations reveal a shape instability that is determined by geometry, where colony eversion occurs for small thicknesses of the contractile shells h and large colony radii R. **d)** Scatter plot of experimental data of the colony radius R as a function of the thickness of the contractile layer h. The data were obtained from 8 colonies from 5 biological replicates and 2 - 4 time points were acquired for each colony. Yellow: colonies that evert 30 min later. Blue: all other colonies. **e)** Channel formation is an essential step for the instability. The time to form a first channel decreases with colony size and increases exponentially with thickness of the contractile ring. Diamonds: time until a first channel is observed in simulations as a function of colony radius. Line is a fit with C_0_/R^2^ (Goodness of fit: R^2^=0.79, RMSE 98.24). Circles: time until a first channel is observed in simulations as a function of the thickness of the contractile layer. Line is a fit with an exponential C_1_exp(C_2_h) (Goodness of fit: R^2^=0.93, RMSE=170.69). **f)** Evolution of channels created with variable radii in the outer contractile shell. Above a critical radius, channels widen, which leads to eversion. Dotted line shows a linear critical radius resulting from theory for a capillary instability.

The process occurs in two stages (Fig. 4a, Movie S12). First, cells from the core of the colony breach through the layer of active cells, forming transient, channel-like structures. Initial breaching and formation of channels are transient and largely reversible processes (Extended Data Figure 14d), with sometimes multiple channels seen simultaneously without changing the roughly spherical shape of the colony on a large scale (Extended Data Figure 12b). However, once a channel starts to grow in width, the spherical geometry becomes unstable, leading to colony eversion.

Simulations reveal that the colony eversion depends on the colony radius R and the thickness of the contractile shell h. An approximately linear function R_c_(h) delineates a regime, R < R_c_(h), where the overall colony shape stays approximately the same, while cells from the core are randomly drifting towards the periphery (Fig. 4c). For R > R_c_(h), colony eversion is mostly observed. The corresponding R-h state diagram obtained from experimental data agrees with the simulation results. During the time-course of the experiments, colony radii and thicknesses of the contractile shells were measured repeatedly. Consistently, the onset of eversion occurred when these variables crossed a critical line separating a stable spheroidal configuration from colony eversion, as predicted by simulations (Fig. 4d).

Since the breaching of the contractile shell is an important step during eversion, we employ simulations to quantify the nucleation of channels (Fig. 4e). The mean first time to form a channel is inversely proportional to the surface area of the colony and depends exponentially on the thickness of the breached layer. Thus, phenomenologically, breaching can be interpreted as resulting from a random walk of the interface hitting the outer surface anywhere. Additional simulations of a simplified two-dimensional system support this interpretation, see Extended Data Figure 13.

Since not all channels immediately lead to eversion, we hypothesized that the instability is non-linear in nature, where perturbations with amplitudes smaller than a critical size decay and only large perturbations trigger global rupture. Such instabilities are characteristic for morphological changes driven by capillarity ^40^ ^41^. To test this hypothesis, we artificially inserted channels of prescribed radius ρ into the outer shell by changing the cell type after the system has mechanically relaxed in the simulations. Consistent with the interpretation of eversion as a non-linear instability, we observed that small channels with ρ < ρ_c_ vanish quickly. Channels with a radius ρ > ρ_c_ grow and eventually lead to eversion (Fig. 4f). The critical radius ρ_c_ depends linearly on the thickness h of the contractile shell in simulations. To explain the instability, we consider an analytical model with an effective surface-tension free-energy and conserved volume (Supplementary Information text). The free energy difference is estimated from the shape of a colony with a channel in the contractile shell and the corresponding reference without a channel. The model shows a metastable free-energy minimum at ρ = 0. A local maximum defines the critical radius ρ_c_, beyond which the channel grows. The model predicts a linear dependence of the critical channel radius on the thickness, ρ_c_ ∼ h, which is clearly satisfied by the simulation data and supports the interpretation of the eversion process to be akin to a capillary instability where the surface tension results from an interplay of from graded activity and cohesion of cells. While channel nucleation can also be facilitated by release of active compression energy from the inside of the colony, simulations of a planar system without compression suggest that this effect is not required (Extended Data Figure 13). Experimentally, it is not possible to visualize the channels of weakly attractive cells within the contractile shell, because it can occur in any direction relative to the imaging plane. It would require three-dimensional characterization of single cell motility which is not possible due to limitations of confocal microscopy in terms of imaging speed and phototoxicity. Instead, we went on to experimentally assess further predictions from this model.

All the evidence we have obtained so far indicates that the molecular motor properties of the T4P govern the morphological remodelling of the gonococcal colonies. To scrutinise this further, we used our model to predict how the system would behave if the attractive interactions increased. As the motor forces considerably exceed the rupture forces, the bacteria undergo continuous tug-of-war within the colonies, and the mechanical properties of the colony are described by the rupture force between the T4P and the binding rate. Simulations predict that eversion occurs less frequently when the probability of T4P binding increases, or when the force scale for T4P detachment increases (see Fig. 5a). Figure 5b illustrates the dynamics of colony with high rupture force and high binding rate where no large-scale inside-out eversion occurs.

**Fig. 5.**
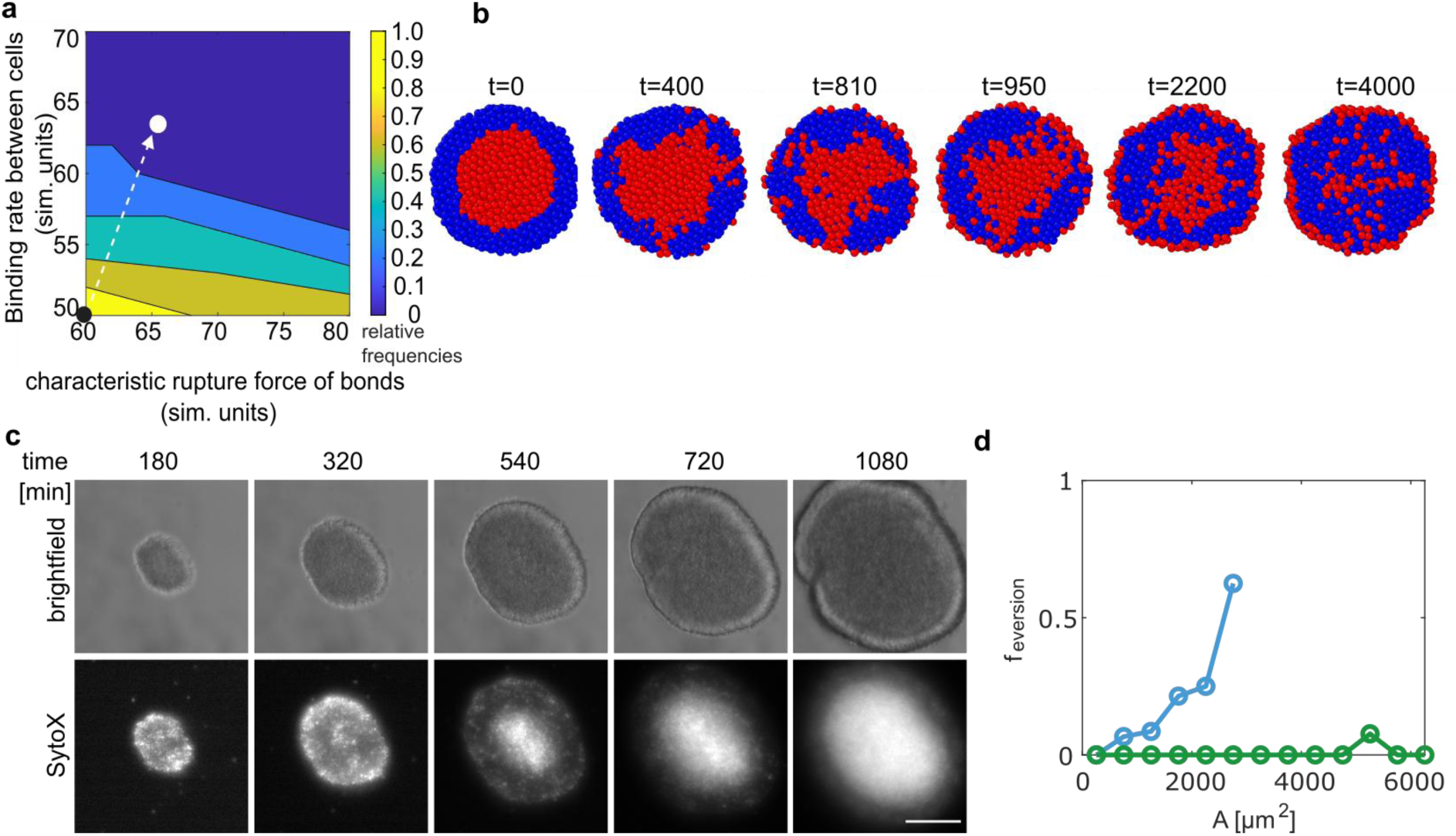
Increasing pilus-pilus attraction inhibits colony eversion. **a)** Simulation results for the relative frequency of eversion as a function of the T4P binding rate and T4P-mediated attractive force. Arrow: predicted positions of wt* (NG150) and *ΔpptA* strain (NG214) in the phase diagram. Colour code: relative frequency. **b)** Time-lapse images of simulated colony dynamics for parameters characteristic of a *ΔpptA* strain. Initially, bacteria with T4P form a contractile shell (blue) around a core of bacteria with reduced T4P activity (red). **c)** Experimentally determined time lapse of colony formed by the *ΔpptA* strain. In each image, the SytoX signal was normalized to 0.2% pixel saturation. Scale bar: 40 µm. The experiment was repeated in 4 biological replicates with a total of N = 39 colonies. **d)** Fraction of everting cells depending on area, A, covered by the colony as determined from brightfield videos. The total number of colonies analysed was N = 568 in 3 biological replicates for wt* (blue) colonies and N = 1158 in 4 biological replicates for *ΔpptA* colonies (green).

To test this prediction experimentally, we examined the dynamics of gonococcal colonies formed by bacteria exhibiting stronger pilus-pilus interactions. Specifically, we used the Δ*pptA* strain, which lacks a phosphotransferase required for posttranslational modification of the pilus ^42^. The loss of posttranslational modification results in higher rupture force and higher binding probability ^31^. Colonies formed by this strain proliferated and accumulated eDNA and dead cells at the centre (Fig. 5c, Movie S11, S12). As predicted, the eversion probability remained very low (Fig. 5d), even when the acquisition time was prolonged to 15 h and the colonies reached a large size.

Overall, our in-silico model reveals the physical mechanism behind eversion, where under spatially inhomogeneous non-equilibrium conditions set by bacterial activity, channel-like structures form that cause a non-linear capillary instability. The model furthermore predicts that the probability of eversion depends on the colony geometry and the strength of pilus-mediated interactions. Experiments confirm these predictions.

### Global colony eversion supports dissemination of bacteria

During eversion, a fraction of cells was shed into the environment (Fig. 1, Fig. 6a). We investigated whether these cells were able to colonize the surface leading to dissemination of bacteria. Shed bacteria generate contrast in the brightfield images (Fig. 6b, Movie S1). To characterize the dispersal of cells from the colonies into the periphery, we measured the standard deviation of the mean of the colony background of these images during the eversion experiment (Fig. 6c). Prior to eversion around 5 h, it remains relatively constant, suggesting cells hardly disperse from young colonies. At the average point of eversion, at 5.5 h, the signal increases abruptly and continues to increase until the end of the experiment, indicating that the surface was covered with a high density of bacteria after colony eversion. The dispersed bacteria formed new colonies (Fig. 6a, b) which underwent twitching motility (Extended Data Figure 10), showing that the bacteria in the colony were alive and capable of T4P retraction. With time the colonies proliferated.

**Fig. 6.**
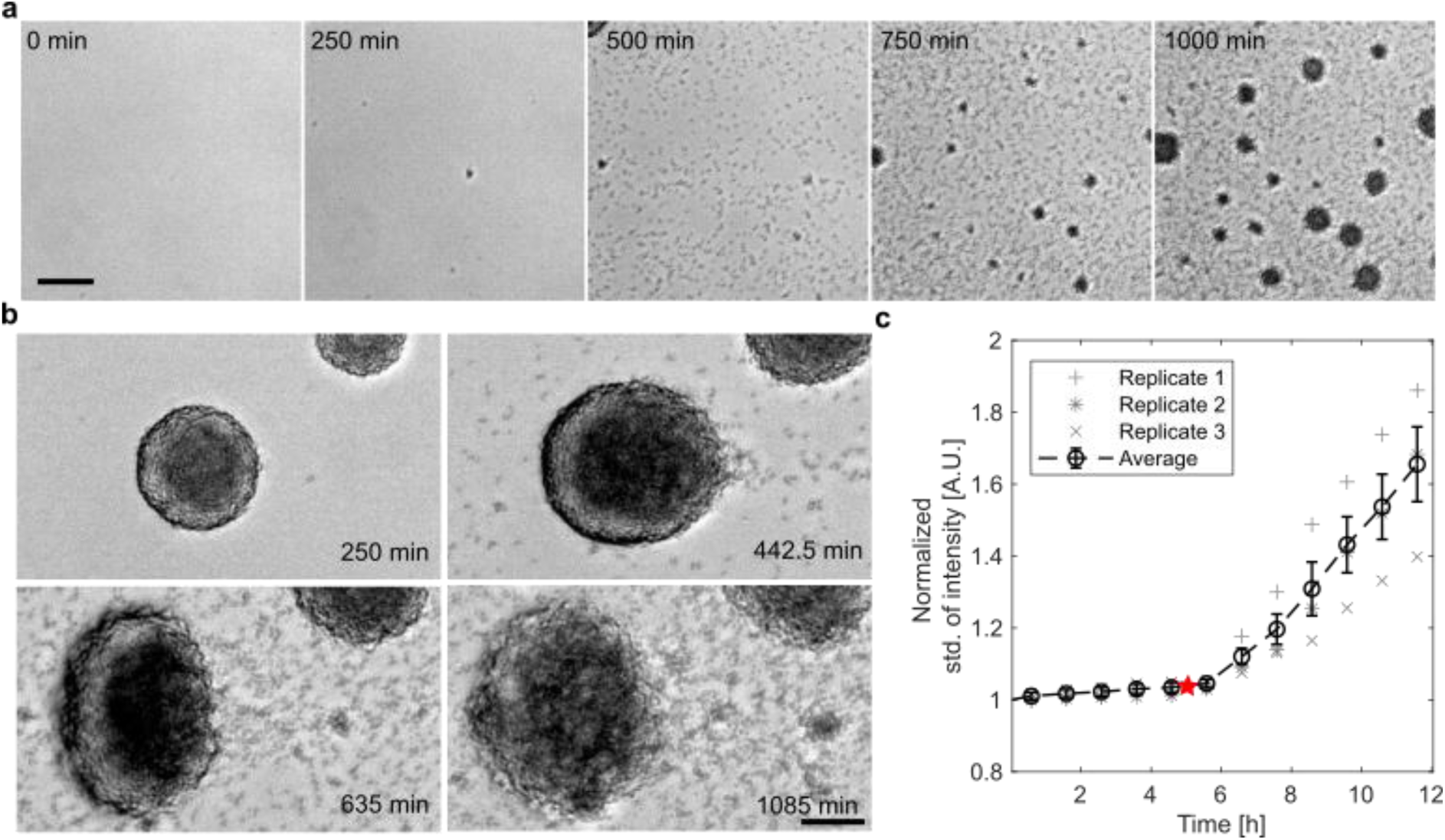
Colony eversion causes dispersal and dissemination. **a)** Brightfield time lapse of area not covered by everting colonies. Strain: NG150. **b)** Typical time lapse of colony during and after eversion. As the colony breaches around 442 min, cells are expelled and form new colonies at later times. Scale bar: 40 µm. **c)** Standard deviation of the background intensity signal. The black dashed line shows the averaged normalized standard deviation of the intensity over 3 biological replicates. Red star: average time point of colony breaching.

Taken together, colony eversion causes dispersal live and dead cells. The living cells actively form new colonies, indicating that eversion enables dissemination of the bacteria. This mechanism of biofilm dispersal differs mechanistically from that observed in biofilms formed by other bacterial systems, including *V. cholerae*, where dispersal was shown to require changes in matrix element expression and specific extracellular matrix degradation ^43^ ^44^ ^45^.

## Discussion

We report a physical mechanism for global restructuring of spherical microbial colonies. Due to limited diffusion of oxygen, a gradient of attractive forces forms that pushes the system out of a quasi-equilibrium state given by the homogeneous spheroidal shape. By combining experimental and simulation approaches, we show that the system can relax towards a lower energy state through a flux of cells from the centre towards the periphery leading to a global eversion of the colony.

This relaxation requires a breaking of the spherical symmetry of the colony. Interestingly, the underlying mechanism differs greatly from bacterial systems studied so far, especially *P. aeruginosa* and *V. cholerae*. Biofilms formed by these rod-like bacteria can be viewed as active nematic systems ^46^, in which orientational instabilities and defect dynamics control large-scale organization and morphogenesis. By contrast, the cell bodies of *N. gonorrhoeae* are almost spherical. Cell division produces dumbbell-shaped complexes, but these do not induce asymmetry because the division planes of successive generations are perpendicular to each other ^47^. Due to surface tension generated by T4P ^15^, gonococcal colonies are spherical. When the attractive forces weaken at the centre, the spherical symmetry nevertheless remains intact, and global re-organisation towards a new stationary state requires a non-linear capillary instability.

*N. gonorrhoeae* has a complex infection and transmission cycle in which oxygen gradients are likely to form. We have demonstrated that the resulting differential attraction between bacteria can generate mesoscopic directed fluxes of bacterial cells. These fluxes enable bacterial dissemination and could potentially promote the spread of infection. In addition to dissemination, cellular flux generates shearing forces that likely affect the extracellular matrix structure and, consequently, the mechanical properties of the colony. Since oxygen heterogeneity is common in bacterial biofilms beyond those formed by the *Neisseriaceae*, it would be interesting to investigate whether it also generates global fluxes for other bacterial species.

## Materials and Methods

### N. gonorrhoeae growth media

To cultivate *N. gonorrhoeae*, bacteria were streaked from a −80°C cryo-stock in GC freeze (milk powder (*Carl Roth GmbH + Co. KG*) in MilliQ water) on gonococcal (GC) base agar plates, containing per 1 L of medium, NaCl 5 g (*Carl Roth GmbH + Co. KG*), K_2_HPO_4_ 4 g (*Carl Roth GmbH + Co. KG*), KH_2_PO_4_ 1 g (*Carl Roth GmbH + Co. KG*), Proteose Peptone No.3 15 g (*Gibco™ Bacto™*), soluble starch 0.5 g (*Sigma Aldrich*), bacto agar 10 g (*Gibco™ Bacto™)* and 10 mL custom made IsoVitaleX (IVX). The IVX was made from D(+)-Glucose 100 g/L (*Carl Roth GmbH + Co. KG*), L-Glutamine 10 g/L (*Carl Roth GmbH + Co. KG*), Cocarboxylase (Thiamine Pyrophosphate) 0.1 g/L (*Sigma Aldrich*), Ferric Nitrate (Fe(No_3_)_3_⋅9H_2_O) 0.02 g/L (*Sigma Aldrich*), Thiamine⋅HCl 0.003 g/L (*Carl Roth GmbH + Co. KG*), 4-Aminobenzoic acid (PABA) 0.013 g/L (*Sigma Aldrich*), β-Nicotinamide adenine dinucleotide (NAD) 0.25 g/L (*Carl Roth GmbH + Co. KG*), Cyanocobalamin (Vitamin B12) 0.01 g/L (*Sigma Aldrich*). IVX was added after autoclaving, and once the medium or agar had cooled down to 55°C. Plates were incubated overnight at 37°C with 5% CO_2_. For liquid media cultivation, GC medium with an identical composition but lacking agar and starch was used. The liquid medium was filtered with a 0.22 µm syringe filter before use.

### Bacterial Strains

All strains used in this study are derivatives of *N. gonorrhoeae* MS11. *N. gonorrhoeae* undergoes pilin antigenic variation, which could influence the outcome of the experiments when investigating T4P interactions. To prevent this, all strains used in this study include a G4 motive deletion, which is required for pilin antigenic variation ^48^. All strains used in this study are listed in Extended Data Table 1.

To construct the *ΔpilE green* strain, the genomic DNA from NG196 *(ΔpilE::cat*) was isolated using a Blood and Tissue Kit (*Qiagen*) and transferred via a spot transformation into NG151 (*iga::P_pilE_ gfpmut3 ermC G4::aac*) ^15^. After selection with chloramphenicol (10 µg/mL final concentration), the resulting deletion of *pilE* was confirmed via PCR using the primers SK5 (CCGCTCGAGCGGTTCCGACCCAATCAACACACC) and SK40 (AATATCTATACTTAAG TCATTTGGCATCAGATGCCTTA). Additionally, the PCR product was sequenced via Sanger sequencing (*Eurofins Genomics*), and the phenotype was confirmed via microscopy. The resulting strain was given the strain number NG335 (Table S1).

### Colony growth and eversion in flow chambers

Bacteria were grown between 15 h and 20 h in flow chamber devices (*Ibidi,* Luer 0.4 mm channel height). The indicated strains were grown on GC+IVX agar plates at 37°C and 5% CO_2_ overnight. Cells were resuspended in fresh GC+IVX liquid medium, and microcolonies were separated by vigorously vortexing the suspension. Next, the OD_600_ of the suspension was determined using a BioPhotometer D30 (*Eppendorf*) and cells were adjusted in GC+IVX to a final OD_600_ of 0.05 in a total volume of 1 mL within a 2 mL reaction tube. Bacterial cells were incubated for 30 min in a shaking-incubator (250 rpm, 37 °C, 5% CO_2_) to form microcolonies.

The tube for the media inflow was placed in a fresh and filtered (0.22 µm) 120 mL GC + IVX medium reservoir. Next, the flow chamber was flushed with the medium by pulling it into the system with a syringe connected to the media outlet tube. Afterwards, this tube was placed in a second reservoir containing a chlorine tablet (*Guest Medical*) to inactivate any incoming bacteria. After 30 min of incubation, the outlet tube of the flow chamber was blocked with a clamp, and 500 µL of the bacterial solution was injected into the flow chamber with a syringe from the outlet side of the flow chamber. The medium reservoir was placed into a 37°C incubator, and the flow chamber was mounted inside a heating chamber at 37°C onto the respective microscope. Media flow was established by connecting the tubing to a peristaltic pump (*model 205U; Watson Marlow*) set to 2.5 rpm, unless otherwise stated.

For image acquisition with single-cell resolution, the *wt_gfp_* strain was used and visualized with an inverted microscope (Ti-E, *Nikon*), equipped with a spinning disk confocal unit (CSU-X1, *Yokogawa*). The excitation wavelength was 488 nm, and a 100x magnification (1.49 NA, oil immersion objective lens, *Nikon*) was used. Image sequences of 4 min with a frame rate of 15 frames/min were taken every 30 min. At the end of each experiment, one additional large brightfield overview picture (2×2) was taken from each colony.

For brightfield images of several colonies and to investigate the emergence of eDNA and dead cells in larger fields of view, imaging was performed using a commercial inverted microscope (Ti Microscope, *Nikon*) with a CMOS camera (C11440-36U ORCA-spark Digital CMOS camera, *Hamamatsu*) equipped with a 20x air Objective. Here, images were taken in 2.5 min or 5 min intervals from three different fields of view. To visualize eDNA and dead cells, 1µL of SytoX (*Invitrogen*) was added to the medium reservoir.

### Growth of *N. gonorrhoeae* in media supplemented with sodium nitrate

To enable *N. gonorrhoeae* to perform denitrification, we supplied 5 mM sodium nitrite (NaNO_2_) (*Carl Roth GmbH + Co. KG)* via the growth medium during the flow chamber experiment. This acts as an electron acceptor in a truncated denitrification pathway of *N. gonorrhoeae* ^49^. Cells were treated as described above, the OD was adjusted to 0.05, and cells were incubated in GC+IVX for 30 min. Afterwards, the microcolonies were transferred to a flow chamber containing GC+IVX+NaNO_2,_ and the colony development was tracked for 15-20 h.

### T4P labeling via maleimide dye

For pili labeling, we used the protocol established by ^34^ with slight adaptation to the respective experimental conditions. A strain *ΔG4 pilE^T126C^* (Ng226), containing a cysteine substitution that can be labeled via a maleimide dye, was used for pilus labeling. Cells were harvested from a plate and adjusted to an OD_600_ of 0.1 in 100 µL cysteine-free retraction assay medium (RAM) and stained with 1 µL of AF488 Alexa Fluor 488 C5 Maleimide (Thermo-Fisher) for 1 h at 250 rpm, 37 °C, 5% CO_2_. Subsequently, cells were centrifuged at 5000g for 3 min, washed with 1.4 mL of GC+IVX, and resuspended again in 120 µL of GC. The sample was split, 60 µL of cell suspension was spotted on a 24 x 24 mm coverslip, and mounted with silicone dots at all four corners of the slip on a microscope slide. The other 60 µL were transferred to a µ-slide 8-well ibiTreat (*Ibidi*). Here, 40 µL of GC were additionally added to cover the entire surface of the µslide with media. In the last step before the imaging, the coverslip microscopy sandwich was sealed with VaLaP (1:1:1, vaseline, lanolin, paraffin). Image acquisition was performed immediately afterwards using the spinning disk confocal microscope.

### Bacterial Growth Curves

To analyze growth behavior of the wt* and the *gfp* - expressing *N. gonorrhoeae* strains, we performed microplate reader-based growth curve experiments. In order to avoid artifacts created by colony formation, *ΔpilE* variants of the strains were used in these experiments. As displayed in Extended Data Figure 11 the expression of *gfp* under the *pilE* promotor, *ΔpilE green,* does not result in a different growth pattern compared to *ΔpilE*. To perform these growth curve experiments, cells were separately harvested from the overnight plates and adjusted to an OD_600_ of 0.1 in triplicate. Next, 10 µL of each was used to inoculate a triplicate of 1mL of GC+IVX in a 24-well plate (*Greiner Bio One*). The plate was incubated in a microplate reader (*TECAN* Infinite M200 plate reader) with a shaking period of 2 min per measurement cycle and 12-point OD measures per well every 10 min at 37°C and 5% CO_2_ for 24 h. The experiment was conducted in three independent replications.

### Micrograph curation and figure preparation

For presentation of micrographs in a figure, representative fields of view were selected in *Fiji* (version 2.16.0/1.54p) ^50^, and the *Enhanced contrast* function was used to normalize the entire time series at 0.2%. If needed, the brightness and contrast of the micrographs were adjusted identically within the compared image sets. To finalize and combine figures, a combination of *Adobe Illustrator* and *Inkscape* were used.

### Image analysis

#### Starting time of eversion, colony radii of brightfield images and direction of eversion

The time points at which colony eversion started, the respective colony radii, and the direction of cellular flow during eversion relative to the media flow were determined from brightfield movies. Starting times were determined manually as the last frame before the colony morphology changed. From these frames, the colony radii were measured by fitting a circle in *Fiji* to the colony, then measuring the area of the circle and derived the radius *r*=√*A*/π. The direction of eversion in Extended Data Figure 3c was manually determined in *Fiji* by measuring the angle from a line drawn from the centre towards the initial direction of the morphological change of the colony during the eversion process, with regard to the flow direction.

#### Fraction of everting colonies as a function of colony radius and colony area

To analyse the fraction of everting colonies as a function of the colony radius, we developed an image analysis pipeline *fractionofeversion.m* in MATLAB to analyse the brightfield videos of everting colonies. Every 20^th^ frame was analysed until the end of the experiment. First, the videos were pre-processed by applying a Gaussian filter and enhancing the contrast. Next, we used the MATLAB function *imfindcircles.m,* which fits a circle to each of the colonies for colonies larger than 10 µm. For each of the images, it was decided manually whether they had already everted or started to evert. This analysis was applied to at least three biological replicates. Finally, the fraction of colonies that have everted for each radius with bins of 5 µm was determined.

For the Δ*pptA* strain we determined the area covered by the colonies in brightfield videos instead of the radii. To filter the noise of the detection we only analysed detected areas above 50 µm^2^. For each of the images we decided manually, whether a colony is everting. From that data we determined the probability of eversion depending on the area covered with a bin size of 500 µm^2^. For comparison, we applied the analysis also on the wt colonies. This analysis was applied to at least three biological replicates.

#### Particle image velocimetry of SytoX fluorescent signal of everting colonies

To investigate the flow of extracellular DNA and dead cells during eversion, we tracked the SytoX fluorescence signal of colonies via the particle image velocimetry app provided for MATLAB ^51^. The videos were pre-processed with *Fiji*. The image processing tool *Enhance contrast* was applied by choosing a pixel saturation of 0.35% for each image in the time stack. We chose the *Multipass Fast Fourier Transform Window Deformation* method for the analysis. The interrogation windows were 64×64 Px to 32×32 Px to 16×16 Px to 8×8 Px and we averaged over 20 frames (100 min) for each colony during the eversion process.

#### Radially averaged effective diffusion constant

To determine the radially averaged effective diffusion constant, the first step was to track single cells from wt*_gfp_* colonies via the *Fiji Trackmate* plug-in. The videos were pre-processed in *Fiji* by adjusting the contrast and applying the *Stackreg* plug-in to exclude global motion of the colony from the movement of single cells. All tracks were saved as .csv data. To analyse the data, a self-developed MATLAB program *RadialMSDwithoutshells.m* was used. The first step was to determine the centre of mass of the colony *C_colony_*, depending on the colony size, this was done automatically by the program, using the .tif file of the colony image sequence. If the colony was too large to fit in the ROI or the analysis was not sufficient, the user could manually choose the centre of the colony by plotting the tracks from the .csv file. The automatic registration was based on the binarised image of the colony and applying the *regionprops.m* MATLAB function which determine the centroid of the colony area. Only tracks with a duration longer than three frames were further processed. The time step was 4 s. Then, from these tracks, the mean-squared displacement (MSD) and the centre of mass of each track, *C_track_*, were calculated. The centre of mass represents the location from which the radial distance to the centre *C_colony_* were determined. We took the MSD of each track after 40 s as a measure of motility and plotted this against the distance of the track of the cell from the centre *C_colony_*. To smooth the data, a moving average over 40% of the length of the data was applied via the MATLAB *movmean.m* function. For the figures we used a moving average over 20% of the length of the data.

We determined the radius of the motile shell with the self-developed MATLAB analysis program *MovingMeanRadialshell.m*. If a motile shell emerged, this was displayed as a peak when the MSD (t = 40s) were plotted against the radial position. Less motile regions were displayed as valleys. Inverting the plot resulted in the valleys being displayed as peaks of the low motility regimes. We determined the peak position via the *findpeak.m* MATLAB function of the low motility regime, also obtaining the information about the half-width of the determined peak. The outer position of the half-width, we defined as the boundary of the highly motile shell. We then subtracted the boundary position from the radius of the colony, which was defined by the maximal distance to centre of the colony. By this, we obtained the radius of the highly motile shell.

#### Analysis of freshly assembled colonies after eversion

We analysed a brightfield area of 253.9 µm x 139.9 μm without colonies present to determine at which time point we observe dispersed cells. To this end, we measured the standard deviation of the mean intensity of this area via *Fiji* and normalised this to the first measured frame of the experiment.

As new colonies were assembled in the space between the everting colonies during and after eversion, we analysed the motility of the freshly assembled colonies with the *Fiji Trackmate* plug-in. The images were pre-processed by inverting the image and applying a Gaussian blur.

#### Mean-squared displacement of single cells in colonies

For the analysis of the motility regimes, we chose three time points per colony: during eversion, 30 min prior eversion and 2-3 h prior to eversion. For each time point, we chose a circular ROI of R = 10 µm and tracked the cell positions via *Trackmate* in *Fiji*. From the cell positions, we determined the MSD with the self-written MATLAB program *DetermineMSDforEversion.m*. We averaged over all analysed colonies and fitted the logarithmically plotted MSD curves linearly within the interval of 50 s to 90 s and obtained the scaling behaviour of the motility regimes.

### Theory

#### Simulation of colonies using Dissipative Particle Dynamics

To simulate the motion of bacterial cells making up the colony, we employ a well-established algorithm similar to dissipative particle dynamics (DPD). The model was previously validated with experimental data ^28^, and is described in detail in the Supplementary Information text.

Briefly, we considered a three-dimensional system containing cells and simulated Newton’s equations of motion including pair-wise repulsive and dissipative forces, forces resulting from thermal fluctuations, and forces resulting from pair-wise attraction of the cells via T4P. The simulation code is was integrated into the molecular dynamics framework of LAMMPS for efficient parallelisation.

In the simulation, the cells are represented as soft spheres of diameter *d*_c_. Each cell can connect to neighbouring cells with 7 pili, modelled as springs that bind and unbind stochastically. The rate constant for bond formation is denoted by *k*_bind_. Once a pilus is bound, this pilus starts to retract, thereby exerting opposite forces *F* on the two connected cells. The maximum retraction velocity of pili is denoted by *v*_0_. Bound pili detach stochastically with a rate *k*_rupt_ *exp*(*F/F*_rupt_), where *k*_rupt_ is a rate constant and, and *F*_rupt_ is a characteristic rupture force of bonds. The simulation parameters give rise to simulation units used in our data representation as follows. For time, we use 1/*k*_rupt_, corresponding to roughly one second. The force scale is *f*_c_, corresponding roughly to 1pN. The length scale is half of the cell diameter, *d*_c_/2. Essential parameters for the results presented in the main text are, for wild-type bacteria fixed as (*k*_bind_ = 50, *v*_0_ = 2, *F*_rupt_= 60). Reduced T4P activity for wild-type bacteria is modelled by using (*k*_bind_ = 5, *v*_0_ = 0.6, *F*_rupt_= 60). For the snapshots of the Δ*pptA* strain in Fig. 5b we use (*k*_bind_ = 65, *v*_0_ = 2, *F*_rupt_ = 70) and for the Δ*pptA* strain with reduced T4P activity (*k*_bind_ = 6.5, *v*_0_ = 0.6, *F*_rupt_= 70). The complete set of parameters is provided in the supplementary text.

#### Data analysis and presentation

All results are presented in simulation units as defined above. The colonies are visualised using the software package OVITO. The flow velocities during eversion presented in Fig. 4b were averaged in a slice with a thickness of 6 cell diameters through the core of the colony. Instantaneous velocities were averaged in a time window of 50 simulation units after onset of the eversion.

## Data availability

The data supporting the findings of this study are available within the Article and its Supplementary Information.

## Supporting information

SI Text Simulations Theory

## Acknowledgments

We thank Gerrit Ansmann and the Maier and Sabass groups for helpful discussions. This work was supported by the Deutsche Forschungsgemeinschaft through grant MA3898 awarded to BM and the Walter-Benjamin grant 545504326 awarded to SW, and the Center for Molecular Medicine Cologne. KZ and BS have received funding from the European Research Council under the European Union’s Horizon 2020 programme, G.a. No. 852585.

## Authors Contributions

SW, IW, MH, KZ, BS, and BM designed research; SW and IW developed, performed, and analysed the experiments; SW performed the majority of the experiments; IW developed the majority of the quantitative image analysis. KZ and BS developed the simulation code, performed data analysis and interpretation. BS and BM supervised the project and acquired funding. SW, IW, KZ, BS, and BM created the figures and wrote the manuscript.

## Movie Captions

**Supplementary Movie 1.** Exemplary time-lapse video showing *N. gonorrhoeae* wt* colonies being incubated in a flow chamber at 37°C with constant flow of GC media. Time-lapse runs for 12 h. Contrast adjusted, Δt = 2.5 min, Scale bar: 40 µm, AVI.

**Supplementary Movie 2.** Exemplary time-lapse video showing *N. gonorrhoeae* wt* colonies being incubated in a flow chamber at 37°C with constant flow of GC media. Time-lapse runs for 16 h. Left brightfield, right SytoX stain. Contrast adjusted, Δt = 5 min, Scale bar: 40 µm, AVI.

**Supplementary Movie 3.** Single Colony of *N. gonorrhoeae wt\** being incubated in a flow chamber at 37°C with constant flow of GC media. Time-lapse runs for 16 h. Left brightfield, right SytoX stain. Contrast adjusted, Δt = 5 min, Scale bar: 40 µm, AVI.

**Supplementary Movie 4.** A combination of three single colonies of *N. gonorrhoeae wt\** out of three independent experiments. Cells have been incubated in a flow chamber at 37°C with constant flow of GC media. Time-lapse runs for 10 h. Left brightfield, right SytoX stain. Contrast adjusted, Δt = 5 min, Scale bar: 40 µm, AVI.

**Supplementary Movie 5.** Exemplary time-lapse video showing *N. gonorrhoeae ΔpilT* colonies being incubated in a flow chamber at 37°C with constant flow of GC media. Time-lapse runs for 16 h. Contrast adjusted, Δt = 5 min, Scale bar: 40 µm, AVI.

**Supplementary Movie 6.** Time-lapse video showing *N. gonorrhoeae* cells expressing the *pilE_cys_* variant covalently labeled with Alexa Fluor 488 C5 maleimide. Indicated on top of the micrographs are the time points after the start of the experiment, as well as the conditions (open: with air-exchange, sealed: air-tight chamber). Contrast adjusted, Δt =0.1 sec, total duration: 5 sec, Scale bar: 2 µm, AVI.

**Supplementary Movie 7.** Exemplary overview time-lapse video of *N. gonorrhoeae wt\** incubated under slow flow conditions at 37°C in constant flow of GC media. Time-lapse runs for 10 h. Left brightfield, right SytoX stain. Contrast adjusted, Δt = 5 min, Scale bar: 40 µm, AVI.

**Supplementary Movie 8.** Exemplary time-lapse video showing a single *N. gonorrhoeae wt\** colony incubated under slow flow conditions at 37°C in constant flow of GC media. Time-lapse runs for 10h. Left brightfield, right SytoX stain. Contrast adjusted, Δt = 5 min, Scale bar: 40 µm, AVI.

**Supplementary Movie 9.** Exemplary time-lapse video showing a confocal plane through the centre of a *N. gonorrhoeae wt green* colony incubated at 37°C in constant flow of GC media. Contrast adjusted, Δt = 4s, Scale bar: 5 µm, AVI.

**Supplementary Movie 10.** Exemplary overview time-lapse video of *N. gonorrhoeae ΔpptA* incubated at 37°C in constant flow of GC media. Time-lapse runs for 18 h. The brightfield images are displayed on the left, the SytoX stain on the right. Contrast adjusted, Δt =5 min, Scale bar 40: µm, AVI.

**Supplementary Movie 11.** Exemplary time-lapse video showing a single *N. gonorrhoeae ΔpptA* colony incubated at 37°C in constant flow of GC media. Time-lapse runs for 18 h. On the left side, the brightfield image is displayed, and on the right side, the SytoX stain. Contrast adjusted, Δt =5 min, Scale bar: 40 µm, AVI.

**Supplementary Movie 12.** Simulation of colony eversion.

**Supplementary Movie 13.** Simulation testing the non-linear nature of the instability by insertion of a channel of defined size into the contractile shell. The radius of colony is *R=*35.78 and the thickness of the contractile shell is *h=*10. The radius of the inserted channel is 2.8. The small channel rapidly closes and a global instability does not occur.

**Supplementary Movie 14.** Simulation testing the non-linear nature of the instability by insertion of a channel of defined size into the contractile shell. The radius of colony is *R=*35.78 and the thickness of the contractile shell is *h=*10. The radius of the inserted channel is 3.6. The channel rapidly widens, which leads to global eversion.

