## Extended Data for "Differential active adhesion and a capillary instability drive global eversion and dissemination of bacterial colonies"

**Extended Data Figures 1-11 and Table 1:**

### Differential active adhesion and a capillary instability drive global eversion and dissemination of bacterial colonies

Stephan Wimmi[1,*], Isabelle Wielert[1,*], Kai Zhou[2,*], Marc Hennes[1], Benedikt Sabass[3,4,§], Berenike Maier[1,§]

1: Institute for Biological Physics, and Center for Molecular Medicine Cologne,

University of Cologne, Cologne, Germany

2: School of General Education, Wenzhou Business College, Wenzhou, China

3: Technical University of Dortmund, Dortmund, Germany

4: Institute for Infectious Diseases and Zoonoses, Department

of Veterinary Sciences, Ludwig-Maximilians-Universitaet Munich, Munich, Germany

* Equal contribution

§ To whom correspondence should be addressed

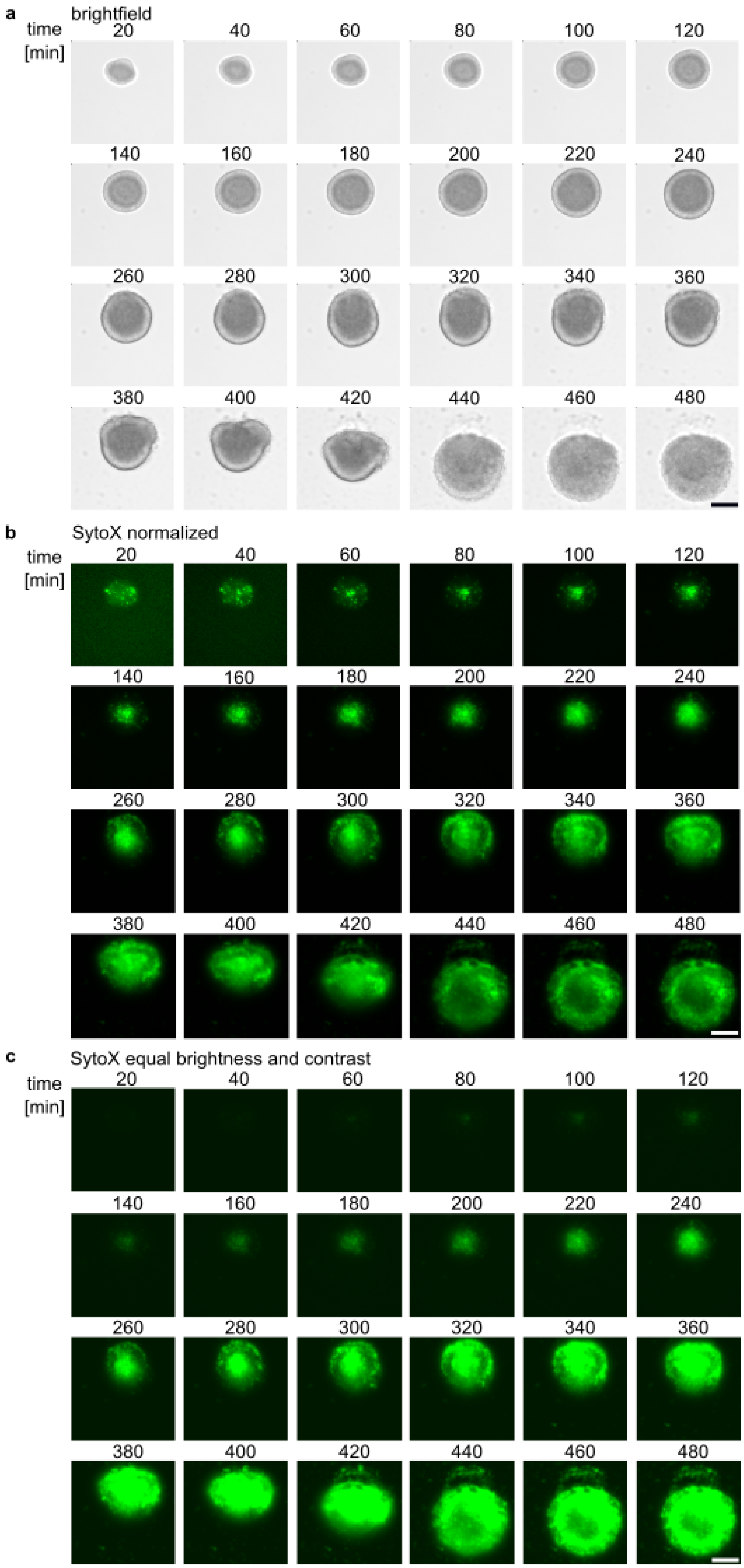
a
brightfield
time [min]
20
40
60
80
100
120
140
160
180
200
220
240
260
280
300
320
340
360
380
400
420
440
460
480
b
SytoX normalized
c
SytoX equal brightness and contrast

**Extended Data Figure 1. Time lapse of colony eversion at higher time resolution. a)** Time course microscopy of *N. gonorrhoeae* colony (NG150) incubated in a flow chamber running at 2.5 rpm at 37°C in brightfield. **b)** SytoX fluorescence signal normalized for representation to 0.2% pixel saturation. **c)** SytoX fluorescence signal with identical and not normalized brightness and contrast values. The elapsed time in minutes is displayed on top of each micrograph; $\Delta t$ = 20 min. Scale bar: 40 µm. This behavior was observed in 7 biologically independent experiments with N = 30 colonies.

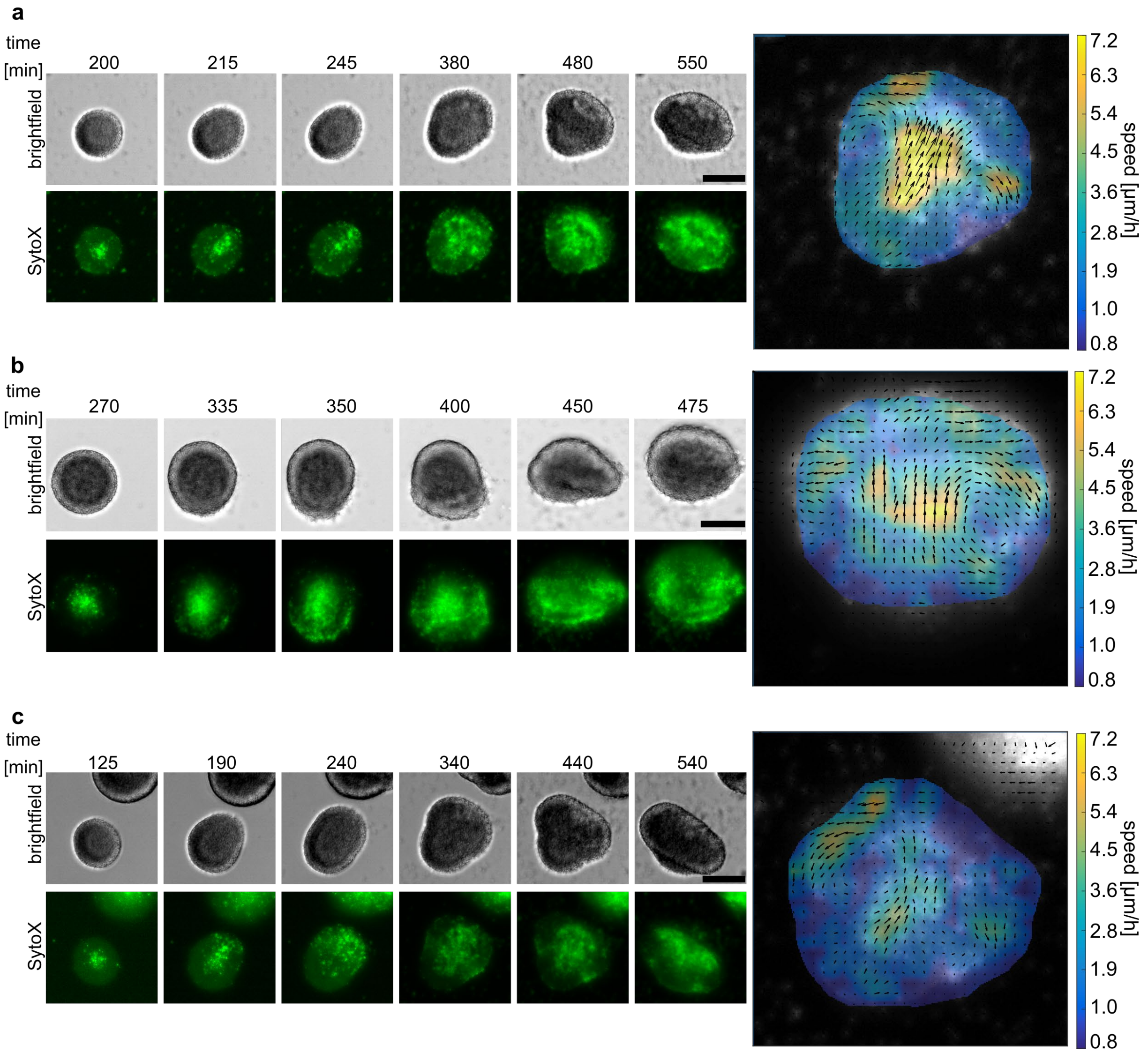


**Extended Data Figure 2. Further examples of everting colonies. a-c)** Three independent sets of representative time-course microscopy experiments with *N. gonorrhoeae* colony (NG150) incubated in a flow chamber running at 2.5 rpm, with brightfield (left, top) and SytoX fluorescence (left, bottom). The elapsed time in minutes is displayed on top of each micrograph. SytoX fluorescence signal normalized for representation to 0.2% pixel saturation. Scale bar: 40 µm. Right: PIV analysis over 100 min of SytoX signal corresponding to the time lapse shown on the left.

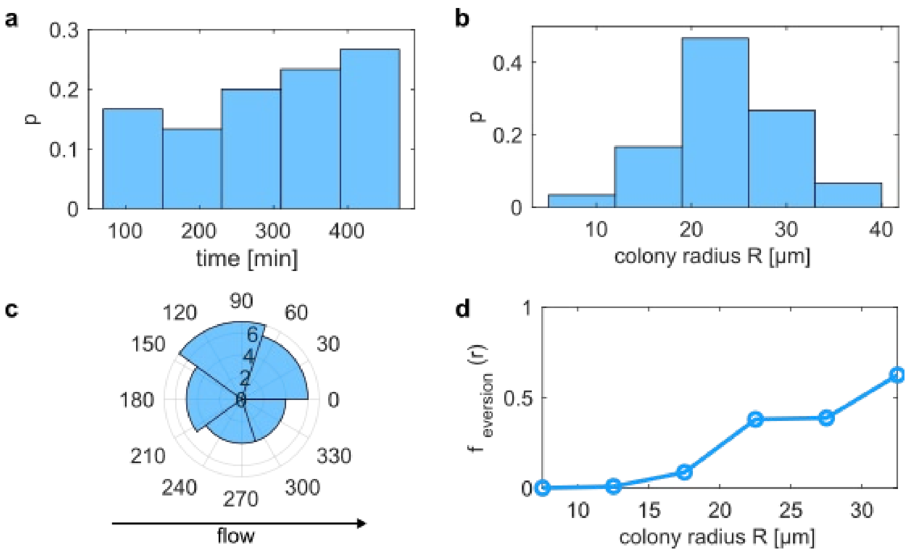


**Extended Data Figure 3. General properties of everting colonies. a)** Relative frequency of times at which eversion starts. N = 30 colonies. Strain: NG150. **b)** Relative frequency of colony radii at start of eversion. N = 30 colonies. **c)** Distribution of angles at which the breaches form relative to the direction of flow. N = 26 colonies. Rayleigh-test indicates random directions of breaching, p = 0.46. **d)** Fraction of everting colonies as a function of colony radius. Images were analyzed at different time points during colony growth. Colony radii were determined and binned. For each bin, the fraction was determined by dividing the number of colonies starting the eversion process by the number of colonies that had not started eversion. N = 2398 colony radii.

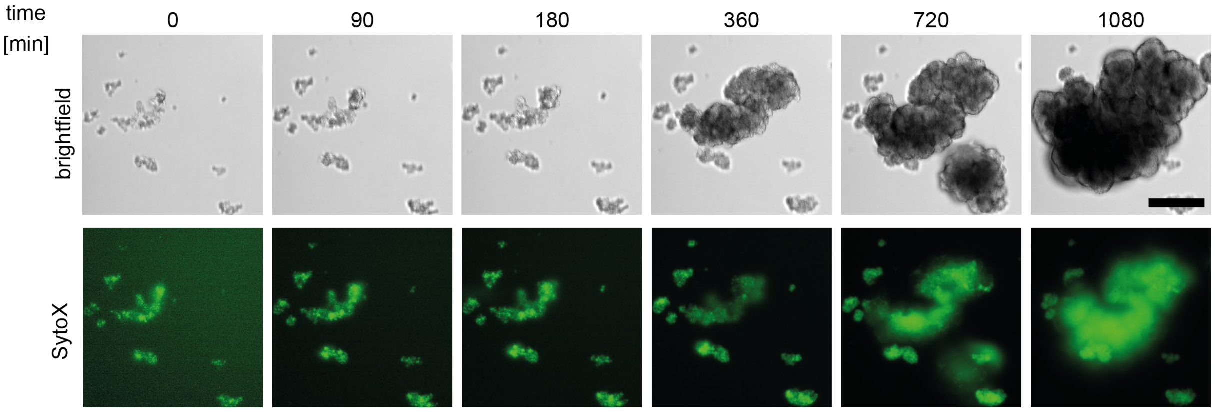


**Extended Data Figure 4. Δ*pilT* strain does not display eversion.** Shown is a typical time lapse of bacteria (NG178) incubated in a flow chamber running at 2.5 rpm at 37°C in brightfield (top) and SytoX fluorescence (bottom) mode. The SytoX fluorescence signal was normalized for representation to 0.2% pixel saturation. The passed time in minutes is indicated on top of the micrographs. Scale bar: 40 µm. The experiment was performed in three biologically independent replicates.

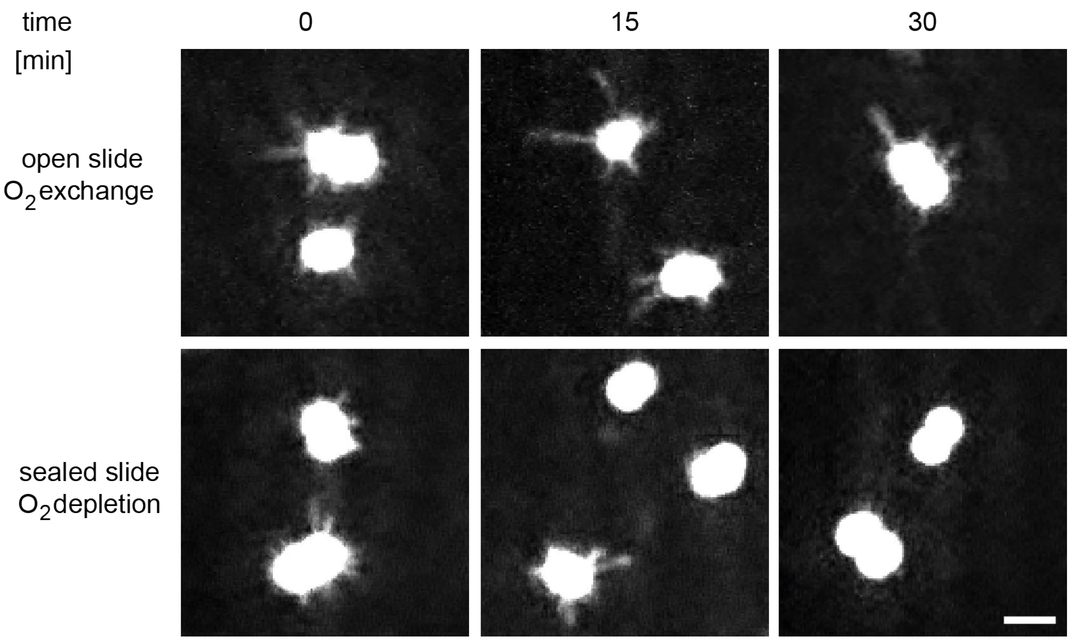


**Extended Data Figure 5. Number of pili decays over time under oxygen depletion.** Representative micrographs of cysteine-maleimide 488-labeled pili (*pilE*$_{cys}$ NG226) corresponding to the quantification displayed in main figure 2. The top row displays micrographs from a sample with oxygen exchange, the bottom row displays micrographs from a sealed slide where oxygen is depleted over time. Elapsed time in minutes is indicated on top of the micrographs. Scale bar: 2 µm. For presentation, the micrographs have been denoised with the *Fiji* plugin *pruedenoise*. The experiment was performed in three biologically independent replicates.

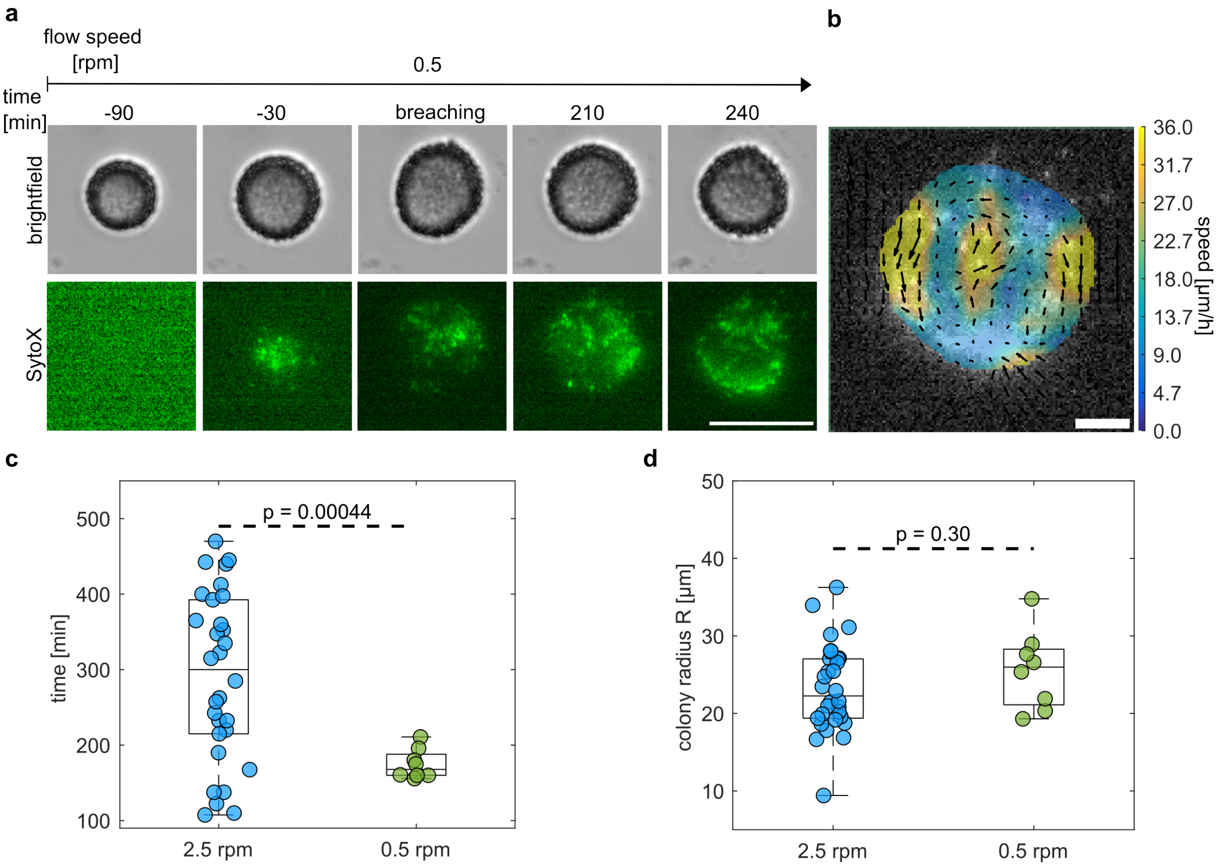


**Extended Data Figure 6. Reduced medium flow results in earlier eversions. a)** Typical time lapse of an evolving colony (NG150) at a flow rate of 0.5 rpm (displayed on top of the micrographs). The elapsed time in minutes and the breaching of the colony's surface are indicated on top of the micrographs. The SytoX signal displayed in these micrographs was normalized to 0.2% pixel saturation. Scale bar: 40 µm. The experiment was performed in three biologically independent replicates. **b)** PIV analysis of the SytoX signal. Scale bar: 10 µm. **c)** Box plots of times at which eversions start for flow speed of 2.5 rpm (blue, N = 30) and 0.5 rpm (green, N = 8). (statistical test: KS-test). **d)** Box plots of colony radii at which eversions start for flow speed of 2.5 rpm (blue, N = 30) and 0.5 rpm (green, N = 8). (statistical test: t-test). Circles: data points of individual colonies. Box plots show the median (central mark), bottom and top edges of the box present 25$^{th}$ and 75$^{th}$ percentiles of the data, respectively.

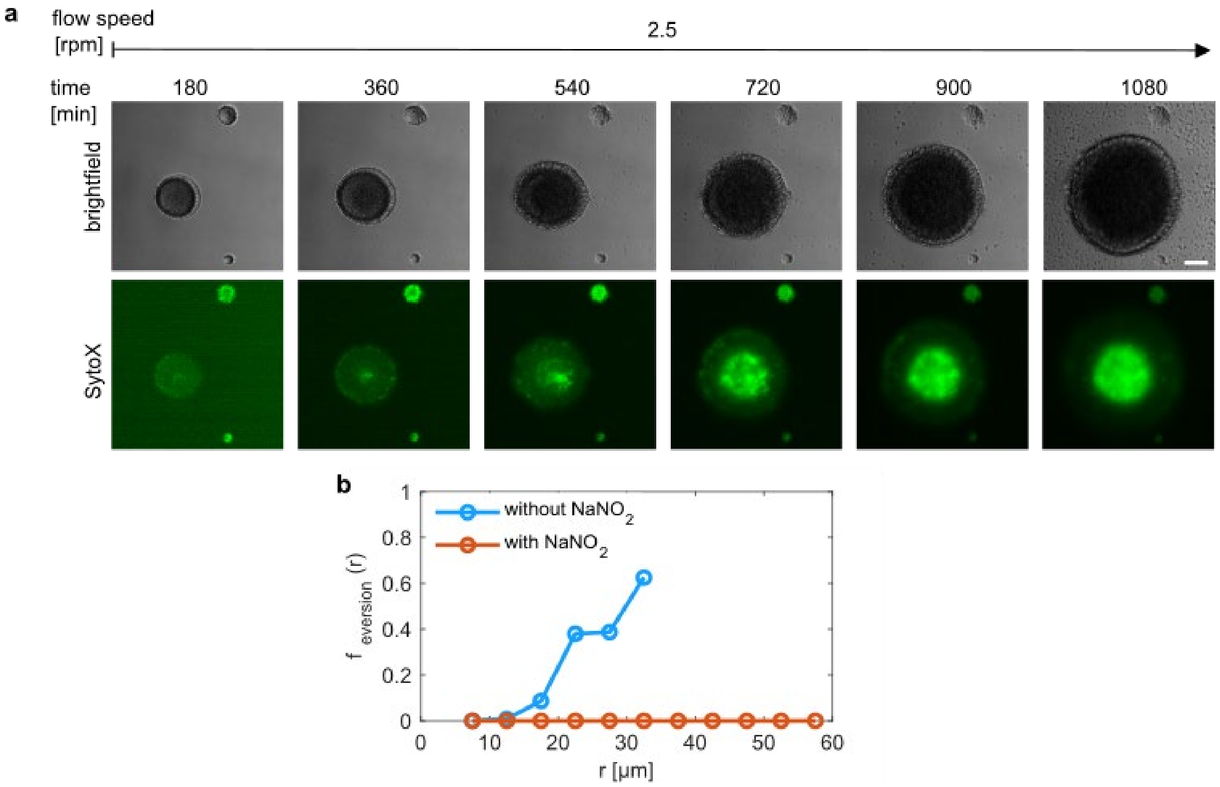


**Extended Data Figure 7. $NaNO_2$ supplementation prevents colony eversion. a)** Representative time lapse microscopy of a growing colony (NG150) at a flow rate of 2.5 rpm (indicated on top) in the presence of 5 mM $NaNO_2$. The experiment was performed in three biologically independent replicates. Top: brightfield, bottom: SytoX fluorescence. The SytoX signal was normalized to 0.2% pixel saturation. **b)** Fraction of everting colonies as a function of colony radii in the presence of 0 mM $NaNO_2$ (blue) (N = 2398 colony radii) and 5 mM $NaNO_2$ (dark orange) (N = 1736 colony radii).

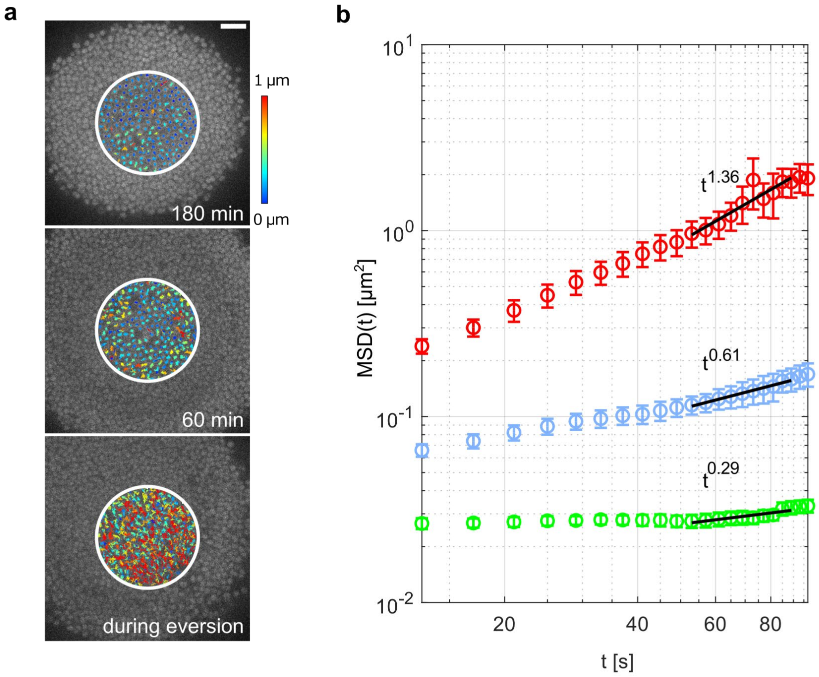


**Extended Data Figure 8. Mean-squared displacement (MSD) analysis shows that the scaling behaviour of the single-cell MSD changes during colony eversion. a)** Overlay of GFP fluorescence (NG151) (grayscale) and trajectories of individual cells tracked (over 4 min) 3 h prior to eversion, 1 h prior to eversion, and during eversion. Scale bar: 5 µm. Color legend: track displacements. **b)** MSD analysis of N ≈ 4000 tracks of cells at the colony centre in a circle of R = 10 µm of 8 colonies 2-3 h prior to eversion (green), 1 h prior to eversion (blue), and during eversion (red) of four biological replicates. Black lines indicate linear fits to determine scaling behavior.

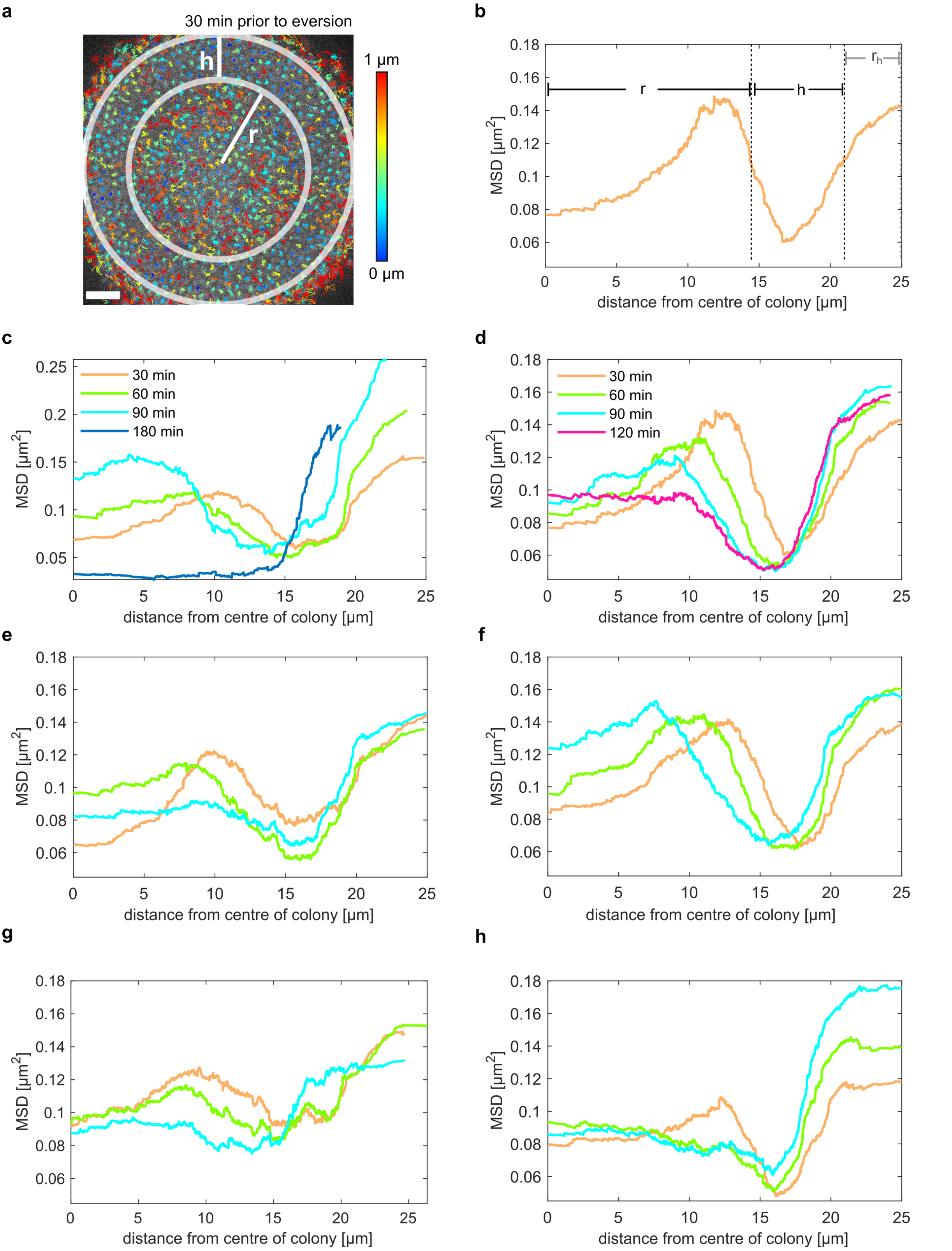


**Extended Data Figure 9. Additional examples for spatially resolved single cell motility. a)** Overlay between GFP fluorescence (NG150) (grayscale) with tracks of cells 30 min prior to eversion. Color legend: track displacements within 4 min. Scale bar: 5 µm. **b)** MSD after 40 s of cells within the colony shown in a). r denotes the distance from the centre to the less motile section of the colony. h is the diameter of the less motile, strongly interacting shell. $r_h$ is the highly motile shell found at the periphery of every colony. In our model, this layer is neglected because it is present at all times and under all conditions. **c)** MSD after 40 s as a function of the distance from the centre of the colony for the time series shown in figure 3 with the hypermotile surface. **d - h)** Further examples of the MSD after 40 s of 5 different colonies.

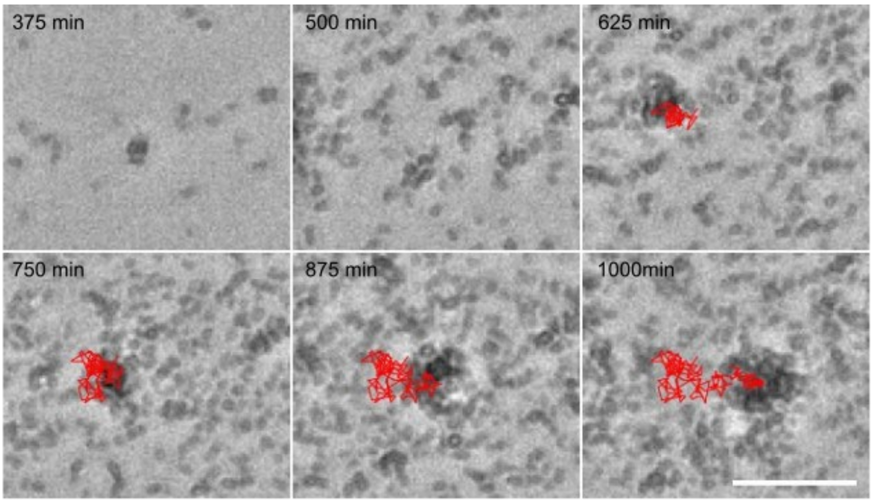


**Extended Data Figure 10. Colony eversion causes dispersal and dissemination**. Brightfield time lapse of newly formed colony (NG150) after eversion of large colonies. The growing colony was tracked in via the *Fiji* plug-in *Trackmate* with the trajectories depicted in red, showing that expelled bacteria form motile colonies. Elapsed time in minutes is indicated in the lower right of each micrograph. Scale bar: 40 µm.

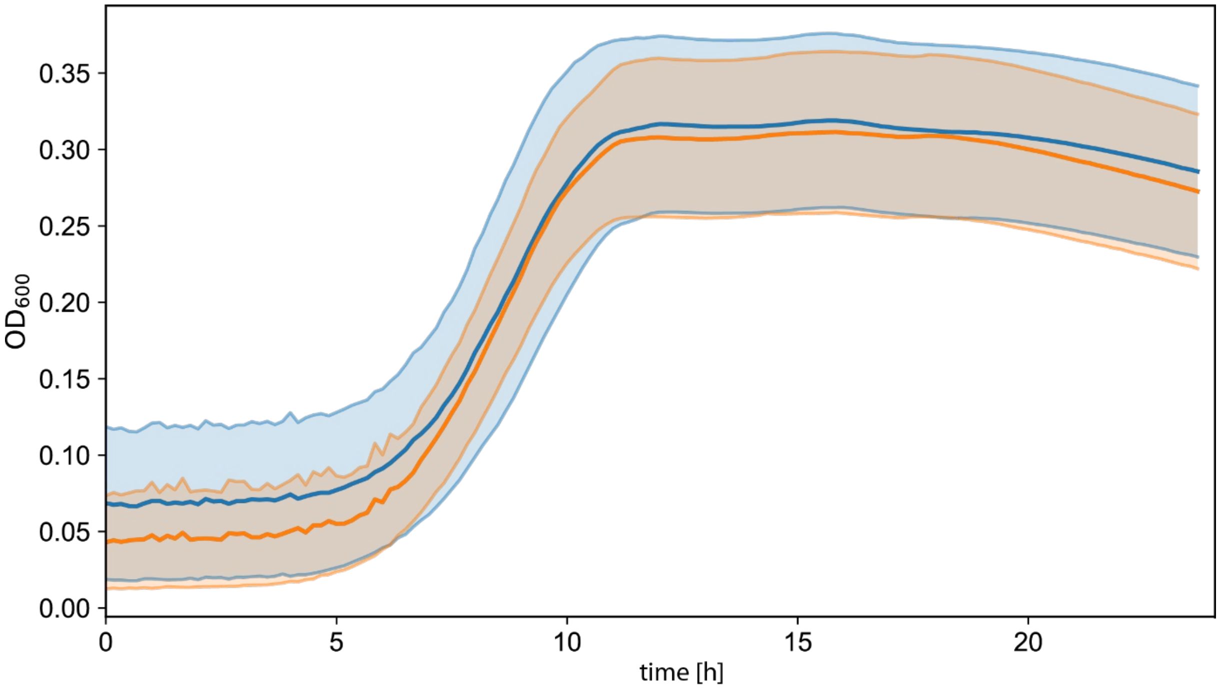


**Extended Data Figure 11. Effect of *gfp* expression on growth.** *N. gonorrhoeae* growth curves were measured at $OD_{600}$ in 24-well plates in the plate reader every 10 minutes. The used strains Δ*pilE* (NG196) (blue) and Δ*pilE green* (NG335) (orange), do not aggregate due to the deletion of the gene encoding the major pilin PilE. Displayed are the average OD600 values from 3 biological replicates; error bands denote the standard deviation between the different measurements.

| Strain | Genotype | Strain number | Reference |
|---|---|---|---|
| *wt** | *G4::apraR* | NG150 | [1] |
| *wt green* | *ΔG4::apraR,*<br>*igA::$P_{pilE}$ gfpmut3 ermC* | NG151 | [2] |
| *ΔpilE* | *ΔG4::apraR, pilE::cat* | Ng196 | [3] |
| *ΔpilE green* | *ΔG4::apraR, pilE::cat,*<br>*igA::$P_{pilE}$ gfpmut3 ermC* | NG335 | this study |
| *ΔpilT* | *ΔG4::apraR pilT::m-Tn3cm* | NG178 | [2] |
| *$pilE_{T126C}$* | *ΔG4::apraR, $pilE_{T126C}$* | NG226 | [4] |
| *ΔpptA* | *ΔG4::apraR, pptA::kanR* | Ng214 | [5] |

**Extended Data Table 1. Strains used in this study**


1 Zöllner, R., Oldewurtel, E. R., Kouzel, N. & Maier, B. Phase and antigenic variation govern competition dynamics through positioning in bacterial colonies. *Sci Rep-Uk* **7** (2017). https://doi.org:ARTN 1215110.1038/s41598-017-12472-7

2 Welker, A. *et al.* Molecular Motors Govern Liquidlike Ordering and Fusion Dynamics of Bacterial Colonies. *Phys Rev Lett* **121** (2018). https://doi.org:ARTN 11810210.1103/PhysRevLett.121.118102

3 Cronenberg, T., Hennes, M., Wielert, I. & Maier, B. Antibiotics modulate attractive interactions in bacterial colonies affecting survivability under combined treatment. *Plos Pathog* **17** (2021). https://doi.org:ARTN e100925110.1371/journal.ppat.1009251

4 Kraus-Roemer, S., Wielert, I., Rathmann, I., Grossbach, J. & Maier, B. External Stresses Affect Gonococcal Type 4 Pilus Dynamics. *Front Microbiol* **13** (2022). https://doi.org:ARTN 83971110.3389/fmicb.2022.839711

5 Zöllner, R. *et al.* Type IV Pilin Post-Translational Modifications Modulate Material Properties of Bacterial Colonies. *Biophys J* **116**, 938-947 (2019). https://doi.org:10.1016/j.bpj.2019.01.020
