## Supplementary material for "Differential active adhesion and a capillary instability drive global eversion and dissemination of bacterial colonies": SI Text Simulations Theory

(Dated: January 11, 2026)

### SIMULATION METHODS

The employed simulation methods are based on established models [1] and we employ an algorithm similar to dissipative particle dynamics (DPD) to simulate the motion of cells inside a colony [2, 3]. In the following, we briefly summarise these methods.

### CELL GEOMETRY AND CONSTRUCTION OF INITIAL STATES FOR COLONY SIMULATIONS

A three-dimensional, Cartesian coordinate system is employed. Individual bacterial cells, also called cocci, are modelled as soft spheres with radius $R$. The position of the center of the cell with index $i$ is denoted by $\mathbf{r}_i$. The vector between a pair of cells with indices $(i, j)$ is denoted by $\mathbf{r}_{ij} = \mathbf{r}_i - \mathbf{r}_j$ and their distance is $r_{ij} = |\mathbf{r}_{ij}|$. Cells repel each other with a potential force modelling volume exclusion. In addition, cells experience dissipative forces resulting from relative motion of neighbouring cells and thermal fluctuations. Each cell has a fixed number of pili. By modelling the pili as dynamic springs that can extend, retract, bind and unbind either with other pili or with the environment, we faithfully represent the stochastic nature of cell-generated forces.

Prior to starting the simulations of the eversion process, the colonies are grown from individual cells through cell division. During the growth phase, each cell divides approximately with rate $\alpha$ into a pair of cells as described in Ref. [1]. In growing colonies, the fraction of dead cells in *N. gonorrhoeae* colonies is reported to be below 5 % [4] and cell death is therefore neglected initially. Once the colonies have reached the desired size to study the eversion process in a given condition, cell growth and division are turned off. After that, the dynamics is governed by passive cell-cell interactions and active retraction of type IV pili as described below.

### CELL-CELL INTERACTIONS THROUGH TYPE IV PILI

Each cell is assumed to have a constant number of pili with the default value of 7, see Tab. I. A pilus of bacterium $i$ is assumed to bind with a rate $k_{\text{bind}}$ to one pilus of a neighbouring bacterium $j$. For binding, the distance between $i$ and $j$, $r_{ij}$, is required to be less than a cutoff distance $d_{\text{bind}}$. This cutoff distance ensures that only bacteria bind to each other when they are in proximity to each other. In some simulations, a distance-based criterion and a Voronoi tessellation are combined to limit pilus interactions only to immediate neighbours that have a distance from each other that is smaller than the cutoff for pilus binding. For simplicity, we assume that pilus-based forces act along the straight lines connecting the centres of cell pairs. It is assumed that two bacteria can only have one pair of pili adhering to each other. Likewise, bundling of pili [5] is also neglected due to its unknown role in cell colonies. Pili of the two cells in a diplococcus do not bind to each other. The pilus-based cell-cell connection is modelled as a spring connecting the centers of two cells. The rest length of the pilus connecting two cells with indices $i$ and $j$ is denoted by $L_{ij}$. The force exerted on the pair of cells is purely attractive and given by

$$f_{ij}^{p} = \min\left[0, -k[r_{ij}(t) - L_{ij}(t)]\right], \tag{1}$$

where $k$ is the pilus' spring constant. Once the pilus is bound, it is assumed to retract. Thus, the effective pilus dynamics employed for our model do not explicitly model non-retracting pili that form passive bonds among cells. The retraction of a pilus leads to a continuous shortening of its rest length as

$$L_{ij}(t) = \max\left[2R, r_{ij}(0) - \int_0^t v_{\mathrm{re}}(t)\mathrm{d}t\right], \tag{2}$$

where $v_{\mathrm{re}}$ is the force-dependent retraction velocity of pili. To describe the force-velocity relationship for pilus retraction motors [6], we employ the linearised relation

$$v_{\mathrm{re}}(t) = \max\left[0, v_{\mathrm{re}}(0)\left(1 - \frac{f_{ij}^{p}}{f_{\mathrm{s}}}\right)\right], \tag{3}$$

where the stall force $f_{\mathrm{s}}$ represents the maximal force a retracting pilus can generate. Furthermore, it is assumed that the bonds between the pili rupture under stress with a force-dependent rate as

$$\gamma_{\mathrm{rupt}} = k_{\mathrm{rupt}} e^{f_{ij}^{p}/F_{\mathrm{rupt}}}, \tag{4}$$

where $k_{\mathrm{rupt}}$ is the pilus rupture rate without loading and $F_{\mathrm{rupt}}$ is a characteristic rupture force. See, e.g., Refs. [5, 7, 8] for related models of pilus dynamics.

### SIMULATED CELLULAR DYNAMICS

Cellular dynamics is governed by Newton's equation of motion with a superposition of different forces, including active forces forces from pili and forces resulting from passive interaction with the cellular environment. For each cell $i$ with mass $m_i$, position $\mathbf{r}_i$, velocity $\mathbf{v}_i$, and force $\mathbf{f}_i$, the equation of motion reads

$$\frac{\mathrm{d}\mathbf{r}_i}{\mathrm{d}t} = \mathbf{v}_i, \qquad m_i\frac{\mathrm{d}\mathbf{v}_i}{\mathrm{d}t} = \mathbf{f}_i. \tag{5}$$

The force acting on each pair of cells consists of conservative forces $\mathbf{F}_{ij}^{c}$, dissipative forces $\mathbf{F}_{ij}^{d}$, thermal fluctuations $\mathbf{F}_{ij}^{r}$ and forces from active pilus retraction $\mathbf{F}_{ij}^{p}$. Overall, the sum of these forces is

$$\mathbf{f}_i = \sum_{j\neq i}(\mathbf{F}_{ij}^{c} + \mathbf{F}_{ij}^{d} + \mathbf{F}_{ij}^{r} + \mathbf{F}_{ij}^{p}). \tag{6}$$

For defining the individual force terms, we employ the vector between the centers of masses $\mathbf{r}_{ij} = \mathbf{r}_i - \mathbf{r}_j$ and the unit vector pointing towards cell $i$ denoted by $\widehat{\mathbf{r}}_{ij} = \mathbf{r}_{ij}/r_{ij}$.

The conservative force acting between pairs of unbound bacteria is

$$\mathbf{F}_{ij}^{c} = \begin{cases} a_0(1 - r_{ij}/d_{\mathrm{con}})\widehat{\mathbf{r}}_{ij} & (r_{ij} < d_{\mathrm{con}}) \\ 0 & (r_{ij} \geq d_{\mathrm{con}}) \end{cases}, \tag{7}$$

where $a_0$ is the maximum conservative force between bacterium $i$ and $j$, the cutoff distance for the repulsive cell-cell interaction is denoted by $d_{\mathrm{con}} = 2R$. For a diplococcus consisting of two spheres, the conservative force due to growth is

$$\mathbf{F}_{ij}^{c} = a_{\mathrm{growth}}(l_i - r_{ij})\widehat{\mathbf{r}}_{ij}, \tag{8}$$

where $a_{\mathrm{growth}}$ is the elastic constant of the spring connecting the two cells of a diplococcus. The dissipative and random forces are, respectively, given by

$$\mathbf{F}^{d}_{ij} = -\gamma\omega^{D}(r_{ij})(\widehat{\mathbf{r}}_{ij}\cdot\mathbf{v}_{ij})\widehat{\mathbf{r}}_{ij}, \tag{9}$$

$$\mathbf{F}^{r}_{ij} = \sqrt{2\gamma k_{\mathrm{B}}T}\omega^{R}(r_{ij})\theta_{ij}\widehat{\mathbf{r}}_{ij}, \tag{10}$$

where $\gamma$ is a friction coefficient, $\omega^{D}$ and $\omega^{R}$ are distance-dependent weight functions, $k_{\mathrm{B}}$ is the Boltzmann constant, $T$ is the ambient temperature and $\theta_{ij} = \theta_{ji}$ is a random number drawn from a Gaussian distribution with zero mean and unit variance. For the distance-dependence of the friction force we choose

$$\omega^{D}(r) = [\omega^{R}(r)]^2 = \begin{cases} (1 - r_{ij}/d_{\mathrm{dpd}})^2 & (r_{ij} < d_{\mathrm{dpd}}) \\ 0 & (r_{ij} \geq d_{\mathrm{dpd}}) \end{cases}, \tag{11}$$

where $d_{\mathrm{dpd}}$ is the cutoff distance for dissipative and random forces. Finally, the forces resulting from retraction of pili are given in their vectorial form by

$$\mathbf{F}^{p}_{ij} = f^{p}_{ij}\widehat{\mathbf{r}}_{ij}. \tag{12}$$

Note that we do not consider the torques generated by T4P between pairs of cells.

### IMPLEMENTATION AND PARAMETER VALUES

The simulation code is integrated into the molecular dynamics simulator LAMMPS [12], which allows an efficient parallelisation while providing great flexibility regarding the model choice. To model the cellular dynamics described above, we wrote a new C++ code. The velocity-Verlet algorithm is used to advance the set of positions, velocities and forces. The code is parallelised and large colonies consisting of thousands of cells can be simulated efficiently. The colonies are visualised with OVITO [13] and the colony images presented here and in the main text represent sections through the center of the tree-dimensional structure. Parameter values that are used internally for the simulations are listed in Tab. I. Whenever alternative parameter values are used, they are provided with the results. Note that the time scale of viscous relaxation is smaller than the time-scale of pilus-based interaction. Thus, inertial effects are negligible. Also, we choose the time scale of the pilus-based interaction to be much smaller than the time-scale of cell division $t_{\mathrm{c}} \ll 1/\alpha$ which occurs during the initialisation phase.

For analysis and presentation of the results, we also chose to fix “simulation scales” that approximately match physically-relevant units. The length scale is the cell radius $d_{\mathrm{c}}/2 \sim 1\,\mu$m. The time scale given by the inverse of the default value of the pilus-unbinding rate-constant $t_{\mathrm{c}} = 1/k_{\mathrm{rupt}} = 1$s, and a force scale of $f_{\mathrm{c}} = 1\,$pN. These scales are listed in Tab. II.

Table I: Choice of parameters in the simulations.

| Parameter | Value | Unit | Reference |
|---|---|---|---|
| cell radius $R$ | 0.5 | $d_\mathrm{c}$ | |
| cell mass $m$ | 0.1 | $f_\mathrm{c} t_\mathrm{c}^2 d_\mathrm{c}^{-1}$ | |
| pilus spring constant $k$ | 500 | $f_\mathrm{c} d_\mathrm{c}^{-1}$ | |
| pilus stall force $f_s$ | 180 | $f_\mathrm{c}$ | |
| maximum pilus retraction speed for normal cells $v_0$ | 2 | $d_\mathrm{c} t_\mathrm{c}^{-1}$ | |
| maximum pilus retraction speed for cells w. reduced activity $v_\mathrm{dead}$ | 0.6 | $d_\mathrm{c} t_\mathrm{c}^{-1}$ | |
| number of pili per cell | 7 | | [9] |
| simulation time step $\Delta t$ | $1 \times 10^{-4}$ | $t_\mathrm{c}$ | |
| division rate $\alpha$ | $1/500$ | $t_\mathrm{c}^{-1}$ | |
| diplococcus growth parameter $\nu_\mathrm{r}$ | 1.0 | $d_\mathrm{c}^{-1}$ | |
| pilus rupture rate $k_\mathrm{rupt}$ | 0.5 | $t_\mathrm{c}^{-1}$ | |
| pilus binding cutoff distance $d_\mathrm{bind}$ | 2.2 | $d_\mathrm{c}$ | |
| pilus binding rate for normal cells $k_\mathrm{bind}$ | 50 | $t_\mathrm{c}^{-1}$ | |
| pilus binding rate for cells w. reduced activity $k_\mathrm{dead}$ | 5 | $t_\mathrm{c}^{-1}$ | |
| pilus-pilus bond rupture force scale $F_\mathrm{rupt}$ | 60 | $f_\mathrm{c}$ | [10, 11] |
| maximum conservative force $a_0$ | 2000 | $f_\mathrm{c} d_\mathrm{c}^{-1}$ | |
| conservative force cutoff $d_\mathrm{con} = 2R$ | 1.0 | $d_\mathrm{c}$ | |
| diplococcus spring constant $a_\mathrm{growth}$ | 2000 | $f_\mathrm{c} d_\mathrm{c}^{-1}$ | |
| friction coefficient $\gamma$ | 300 | $f_\mathrm{c} t_\mathrm{c} d_\mathrm{c}^{-1}$ | |
| thermal energy scale $k_\mathrm{B}T$ | 0.02 | $f_\mathrm{c} d_\mathrm{c}$ | |
| dissipative and random force cutoff $d_\mathrm{dpd}$ | 1.7 | $d_\mathrm{c}$ | |

Table II: Conversion of internal scales to simulation units for data analysis.

| Simulation unit | internal scale in simulations | physical value |
|---|---|---|
| length scale | $0.5\, d_\mathrm{c}$ | $\sim 1\,\mu\mathrm{m}$ |
| time scale | $1/k_\mathrm{rupt}$ | $\sim 1\,\mathrm{s}$ |
| force scale | $f_\mathrm{c}$ | $\sim 1\,\mathrm{pN}$ |

### ANALYSIS OF THE SIMULATION RESULTS

### EVERSION ONLY OCCURS FOR ACTIVELY CONTRACTING CELLS IN SIMULATIONS

To simulate the reduced activity of type IV pili in the idealised representation, the maximum retraction velocity $v_0$ of the pili and their binding rate constant are reduced in the core region with a predefined radius, once the colony has reached a stationary state. Since the core of the colony presumably maintains some level of pilus activity during the experiment, the maximum retraction speed is not set to zero in the simulated colony core, but is reduced to less than half its value in the contractile shell. We also performed numerical experiments where we simply set $v_0 = 0$ in the core and thus assumed a completely passive colony centre. Such parameter values result in a rigid core that suppresses eversion in simulations. To investigate how much the eversion process relies on pilus activity, we vary the maximum retraction velocity in both the outer and inner regions. This reduction of the pilus activity decreases the likelihood of observing an eversion event in simulations. Extended Data Figure 12 depicts representative snapshots from simulations for illustration. We fix a colony geometry for which eversion is observed using our default simulation parameters, see Extended Data Fig. 12a, and then systematically reduce $v_0$ in both the active shell and the passive core. While channel formation can still be observed for reduced activity, global colony eversion does not occur, see Extended Data Fig. 12b. When the maximum retraction velocity is reduced further, we only observe a diffusive passage of passive cells through the contractile shell, see Extended Data Fig. 12c. Consequently, a reduction of pilus activity prevents global eversion and the passive cells from the core can still accumulate on the periphery of the colony.

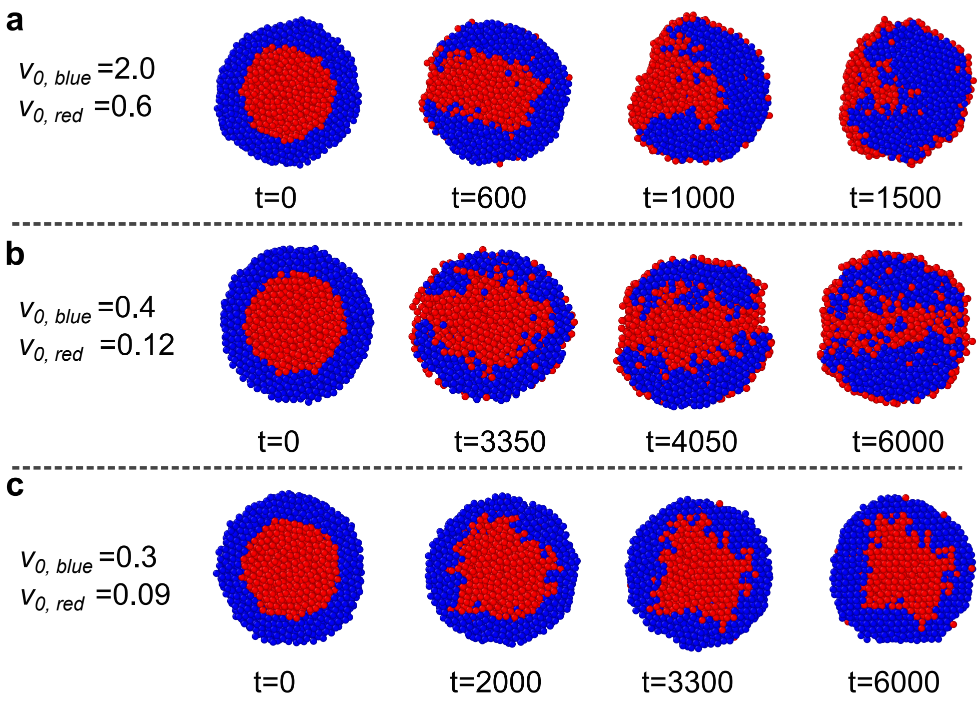


Extended Data Figure 12: Eversion is driven by active retraction of pili. (a) Snapshots of the eversion process observed for cells that are retracting their pili with the default maximum velocity $v_0$. (b) Snapshots of the colony evolution for reduced retraction velocity. In spite of channel formation, a rapid, large-scale folding of the structure is not visible. (c) Snapshots of the colony evolution for strongly reduced retraction velocity. The more passive bacteria diffuse to the periphery of the colony and this is a gradual process without large-scale eversion. Default simulation units are used, see Tab. I.

### FORMATION OF CHANNELS AND INSTABILITY IN A PLANAR GEOMETRY

The spherical shape of colonies introduces a length scale and some extra physical complexity. In interface physics, one can frequently study local phenomena by approximating the large-scale shape through a planar setup. Therefore, we start by studying a planar geometry where we place a contractile layer of cells above a thick layer of passive cells. Extended Data Figure 13 shows results from simulations of the instability in a planar geometry where we chose periodic boundary conditions. We find that the dynamics of channel formation are similar in this geometry and in the spherical geometry. The time to form a first channel $\tau$ depends on the surface area $A$ as $1/\tau \propto A + A_0$ with $A_0$ being a constant. Hence, the rate of channel formation per area approaches a constant for large $A$. Furthermore the time to form a first channel depends exponentially on the thickness of the layer of active bacteria $h$ as $\tau \propto e^{z_0 h}$, with $z_0$ being a positive constant. Overall, the data from simulations of different conditions is fit well by a function $1/\tau = c_1 e^{-z_0 h}(A + A_0)$, with $c_1$ being a second constant.

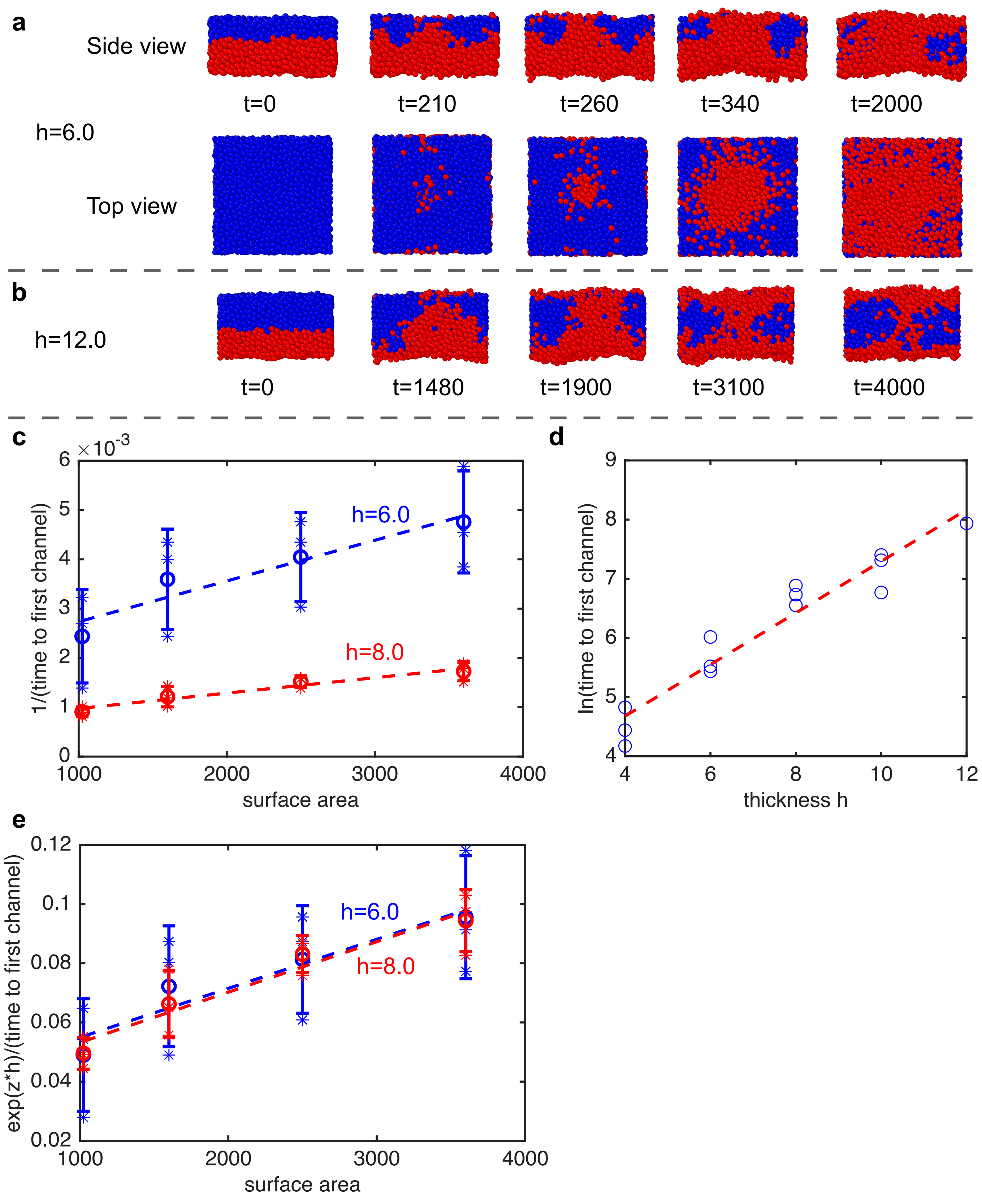
a
Side view
h=6.0
Top view
t=0
t=210
t=260
t=340
t=2000
b
h=12.0
t=0
t=1480
t=1900
t=3100
t=4000
c
×10^-3
1/(time to first channel)
h=6.0
h=8.0
surface area
d
ln(time to first channel)
thickness h
e
exp(z*h)/(time to first channel)
h=6.0
h=8.0
surface area

Extended Data Figure 13: Simulations of a channel formation in a planar-periodic geometry. Parameters for simulation of bacteria were chosen as for the simulations of spherical colonies. (a) Channel formation and spatial reorganisation of active (blue) and less active (red) bacteria. The thickness of the active bacteria is $h = 6$. (b) For a thicker layer of active bacteria, $h = 12$, the channel appears at a later time. (c) The inverse time to form the first channel is proportional to the surface area, with a proportionality factor depending on the thickness of the active layer $h$. (d) The time to form the first channel $\tau$ depends exponentially on the thickness of the active layer as $\tau \propto e^{z_0 h}$, with $z_0 \simeq 0.5$. (e) The data shown in (c) collapses to one curve after rescaling the time to form the first channel with its exponential dependence on $h$. Default simulation units are used in all plots, see Tab. I.

### SIMULATION RESULTS FOR SPHERICAL COLONIES

Spherical colonies can build up internal pressure and the compression of the passive core is clearly measurable. To find out whether the energy stored in compression changes the results significantly, we conduct extensive simulations varying the colony size parameters. Extended Data Figure 14 shows additional results that complement data shown in the main text. For spherical colonies, the time until formation of the first channel also depends exponentially on the thickness of the active layer $h$, $\tau \propto e^{z h}$, with a factor $z$ that depends slightly on the size of the overall colony, see Extended Data Fig. 14b. The factor $z$ deviates from its value in the planar configuration, $z_0$, most notably for small colonies and large $h$. However, Extended Data Fig. 14b also illustrates that different prefactors for spherical colonies do not differ drastically from the value $z_0$ measured in the planar setup. Thus, the internal pressure built-up by the contractile shell around spherical colonies does not affect the dynamics of initial channel formation qualitatively.

Formation of the first channel frequently does not directly trigger the instability. Extended Data Figure (14)d shows the delay time between formation of a first channel and the formation of a channel that actually triggers eversion. The plot suggests that larger colony radii reduce the delay time.

To quantify the nature of the instability, channels are produced in a controlled fashion by changing the cell type in a defined volume in the contractile shell after an initial relaxation phase, as also explained in the main text, see Extended Data Fig. 14e. We find that, similar to classical capillary instabilities [14, 15], the radius of the channel determines whether the channel will shrink or grow. The critical radius of the channel, $\rho_c$, above which eversion occurs, depends linearly on the thickness of the contractile layer $h$ and the slope of the function slightly depends on the Radius $R$, see Extended Data Fig. 14f-g. The function $\rho_c \simeq h(C_1 + C_2/R^2)$ with two constants $C_1$ and $C_2$ fits the simulation data well and suggests that the radius dependence amounts to a secondary correction to the result for the planar system for the simulated cases. Note that the form of this correction function implies the existence of an additional length scale in the problem, beyond the two scales $h$ and $R$, which are fixed by geometry and are already part of this formula. It is not apparent how such a scale would emerge from a more detailed analysis of the problem in any model built only on minimal-surface arguments.

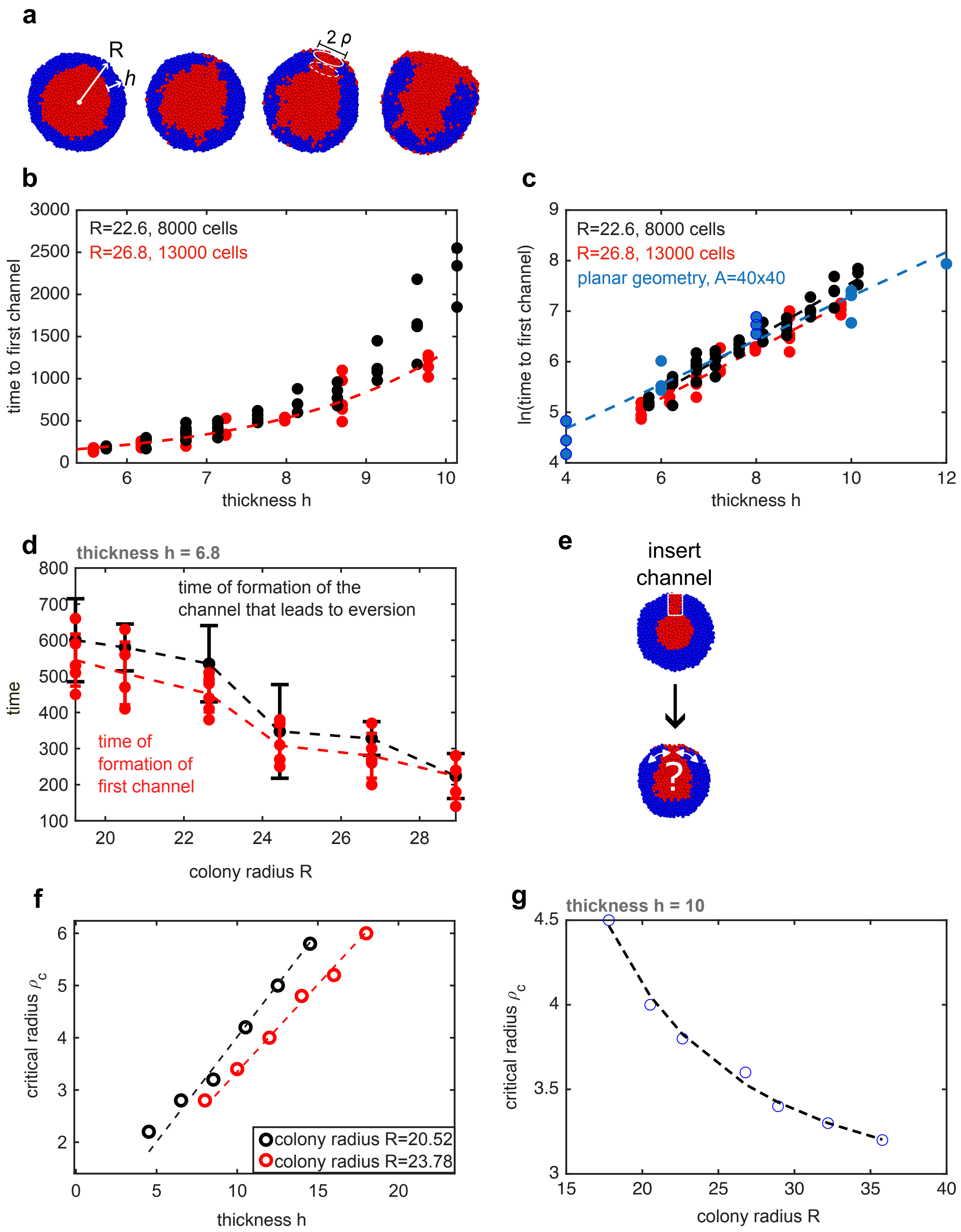

a
R
h
2 ρ
b
R=22.6, 8000 cells
R=26.8, 13000 cells
time to first channel
thickness h
c
R=22.6, 8000 cells
R=26.8, 13000 cells
planar geometry, A=40x40
ln(time to first channel)
thickness h
d
thickness h = 6.8
time of formation of the
channel that leads to eversion
time of
formation of
first channel
time
colony radius R
e
insert
channel
?
f
critical radius $\rho_c$
colony radius R=20.52
colony radius R=23.78
thickness h
g
thickness h = 10
critical radius $\rho_c$
colony radius R

Extended Data Figure 14: Simulation of eversion in a spherical geometry. (a) Simulation snapshots. (b) Time until formation of the first channel vs. thickness of the contractile shell $h$ for colonies of two sizes. (c) The time until first channel formation scales as $\tau \propto e^{z\,h}$, with a time scale $z$ that depends slightly on the size of the overall colony. The scale $z$ deviates from its value in the planar configuration, $z_0$, for small colonies. (d) To assess reversibility of channel formation, we quantify the times of first channel formation and the time of formation of the channel that eventually leads to eversion. Typically, multiple channels are formed before an eversion occurs. However, for very large colonies, the first channel frequently leads to eversion. (e) To quantify the nature of the instability, channels are produced in a controlled fashion by changing the cell type in a defined volume in the contractile shell after an initial relaxation phase. (f) The critical radius of the channel, above which eversion occurs, depends linearly on the thickness of the contractile layer. (g) For a fixed thickness $h$, the critical radius $\rho_\mathrm{c}$ decreases with larger colony sizes. The dashed line shows a fit with the function $\rho_\mathrm{c} = h(C_1 + C_2/R^2)$. All plots are represented in our default simulation units, see Tab. I.

### THEORY FOR CAPILLARY INSTABILITY OF SPHERICAL BACTERIAL COLONIES

To model the mechanical instability after the formation of a channel, we consider an idealised setup consisting of two concentric spheres. The surface tension between the active shell and the surrounding fluid is denoted by $\gamma_{\mathrm{AV}}$. The surface tension between the active shell and the passive core is denoted by $\gamma_{PA}$. The surface tension between the passive bacteria and the surrounding fluid is denoted by $\gamma_{\mathrm{PV}}$. The free energy of the reference state results from the surface tension of the concentric configuration and is given by

$$\mathcal{F}_0 = \gamma_{\mathrm{AV}} 4\pi R^2 + \gamma_{PA} 4\pi (R - h_0)^2, \tag{13}$$

where $R$ is the constant outer radius of the colony and $h_0$ is the initial thickness of the contractile shell. Now suppose that a channel filled with passive bacteria appears in the contractile shell. For simplicity, this channel is modelled as a cylinder with radius $\rho$. The thickness of the contractile shell in the new configuration is denoted by $h$ where $h \neq h_0$ due to incompressibility. The free energy of the configuration with the channel is given by

$$\mathcal{F}_1 = \gamma_{\mathrm{AV}} \left(4\pi R^2 - \pi\rho^2\right) + \gamma_{\mathrm{PV}} \pi\rho^2 + \gamma_{\mathrm{PA}} \left(4\pi (R-h)^2 - \pi\rho^2 + 2\pi\rho\, h\right), \tag{14}$$

where we assumed $\rho \ll R$ to simplify the expression. The free energy difference reads

$$\Delta\mathcal{F} = \mathcal{F}_1 - \mathcal{F}_0. \tag{15}$$

We determine $h$ from volume conservation for the contractile shell

$$\frac{4}{3}\pi R^3 - \frac{4}{3}\pi (R - h_0)^3 = \frac{4}{3}\pi R^3 - \frac{4}{3}\pi (R-h)^3 - \pi\rho^2 h, \tag{16}$$

where the left hand side is the volume in the original concentric-sphere configuration and the right hand side is the volume in the configuration with a channel. Insertion of the resulting equation for $h$ into Eq. (15)

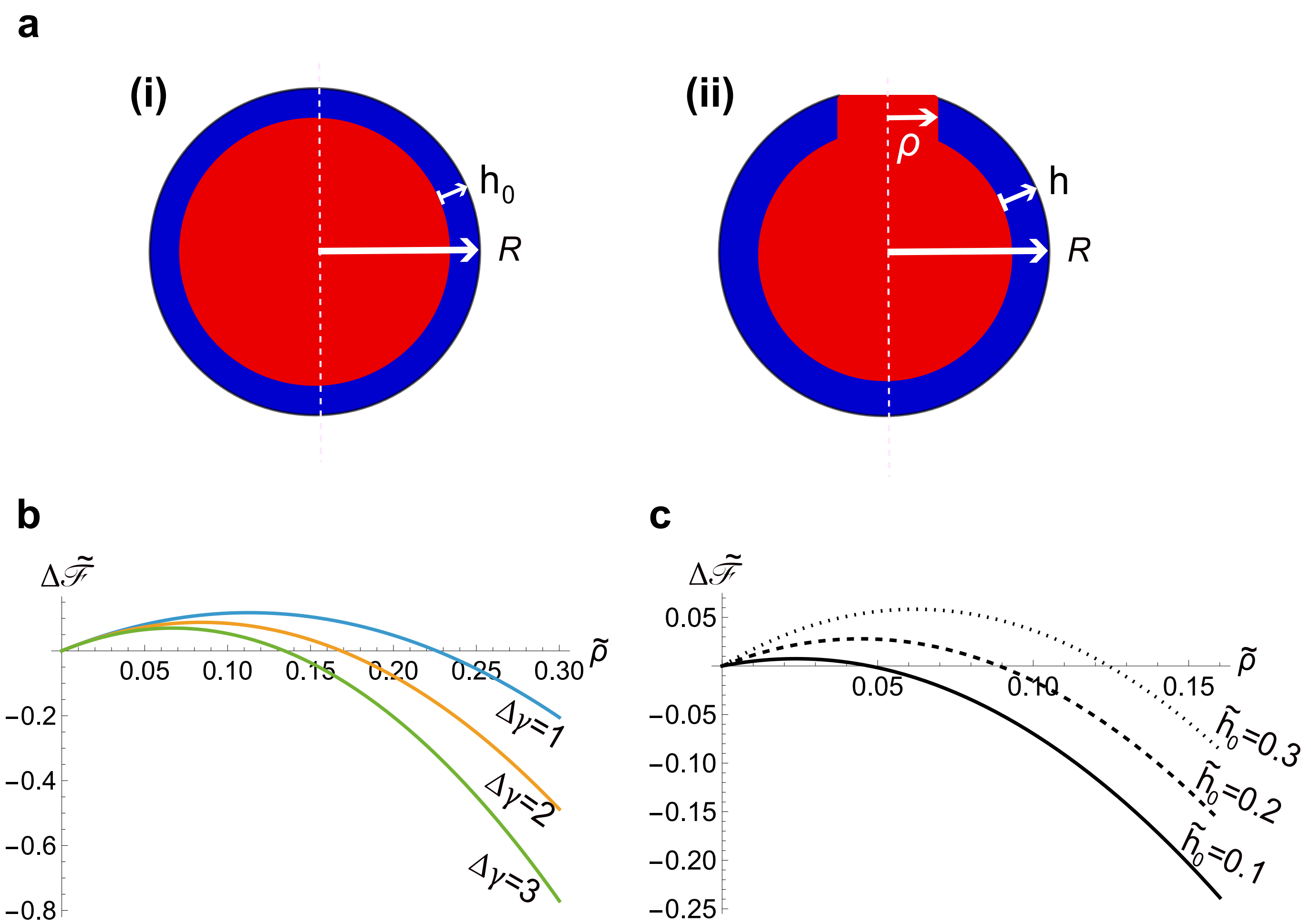


Extended Data Figure 15: An idealised model representing the colony instability. (a) Sketch of the model with (i) the reference state with radius $R$ and thickness of contractile shell $h_0$ and (ii) the configuration with a channel of radius $\rho$. (b) Free energy difference $\Delta\tilde{\mathcal{F}} = \Delta\mathcal{F}/(\gamma_{\rm PA}\,\pi R^2)$ as a function of $\tilde{\rho} = \rho/R$ for different values of the relative surface tension difference $\Delta\gamma$. (c) b) Free energy difference $\Delta\tilde{\mathcal{F}}$ as a function of $\rho/R$ for different thicknesses of the contractile layer $\tilde{h}_0 = h_0/R$.

yields a lengthy expression for the free energy. An expansion for large $R$ produces

$$\frac{\Delta\mathcal{F}}{\gamma_{\rm PA}\,\pi} \approx 2h_0\rho - (\Delta\gamma + 1)\rho^2 - \frac{2h_0\rho^2}{R} + \frac{(h_0\rho^3 - 4h_0^2\rho^2)}{2R^2}, \tag{17}$$

where we introduced the relative surface tension difference $\Delta\gamma \equiv (\gamma_{\rm AV} - \gamma_{\rm PV})/\gamma_{\rm PA}$. Extended Data Figure 15 shows exemplary full solutions of the non-dimensional free energy $\Delta\tilde{\mathcal{F}} = \Delta\mathcal{F}/(\gamma_{\rm PA}\,\pi R^2)$ as a function of $\tilde{\rho} = \rho/R$. The function has a local minimum at $\rho = 0$ and a maximum at $\rho = \rho_{\rm c} > 0$. Thus, the homogeneous solution without a channel, $\rho = 0$, is only locally stable, a hallmark of capillary instabilities [14, 15]. The free energy can be lowered by the formation of a large enough channel $\rho > \rho_{\rm c}$. The critical channel radius is determined by $\frac{\partial \Delta\mathcal{F}}{\partial\rho} = 0$. In the limit $R \to \infty$, where the equations become independent of $R$, we find

$$\rho_{\rm c} \approx \frac{\gamma_{\rm PA} h_0}{\gamma_{\rm AV} - \gamma_{\rm PV} + \gamma_{\rm PA}} = \frac{h_0}{\Delta\gamma + 1}, \tag{18}$$

which explains why in Extended Data Fig. 14f the critical channel radius is proportional to $h_0$. We also note that the slope of the critical line is only determined by the surface tension difference $\Delta\gamma$. Higher-order correction terms in powers of $h_0/R$ can be easily calculated. However, the scaling $\rho_\mathrm{c} \simeq h(C_1 + C_2/R^2)$ found in the simulation data suggests the existence of an additional length scale at order $1/R^2$. Therefore, the simple surface-tension model becomes insufficient at that order. Note that also a more sophisticated quasi-equilibrium solution, where the shape of the inner surface is calculated under the assumption of instantaneous relaxation, does not provide the additional length scale and is therefore of little value for the present problem. However, model extensions such as a consideration of elastic deformations that cause effective bending and compression modes would result in additional scales, as would a consideration of the finite size of bacteria. Thus, the higher-order behaviour of the threshold for the capillary instability remains an interesting open question for future studies.

---

*
